# Matrix Gla Protein acts as a driver of stemness and tumor initiation in ovarian cancer

**DOI:** 10.1101/2022.12.06.519263

**Authors:** V. Nieddu, V. Melocchi, C. Battistini, G. Franciosa, M. Lupia, C. Stellato, G. Bertalot, J.V. Olsen, N. Colombo, F. Bianchi, U. Cavallaro

## Abstract

Ovarian cancer (OC) displays the highest mortality among gynecological tumors, mainly due to early peritoneal dissemination, the high frequency of tumor relapse following primary debulking and the development of chemoresistance. All these events are thought to be initiated and sustained by a subpopulation of neoplastic cells, termed ovarian cancer stem cells (OCSC), that are endowed with self-renewing and tumor-initiating properties. This implies that interfering with OCSC function should offer novel therapeutic perspectives to defeat OC progression. To this aim, a better understanding of the molecular and functional makeup of OCSC in clinically relevant model systems is essential.

We have profiled the transcriptome of OCSC vs. their bulk cell counterpart from a panel of patient-derived OC cell cultures. This revealed that Matrix Gla Protein (MGP), classically known as a calcification-preventing factor in cartilage and blood vessels, is markedly enriched in OCSC. Functional assays showed that MGP confers several stemness-associated traits to OC cells, including a transcriptional reprogramming. Patient-derived organotypic cultures pointed to the peritoneal microenvironment as a major inducer of MGP expression in OC cells. Furthermore, MGP was found to be necessary and sufficient for tumor initiation in OC mouse models, by shortening tumor latency and increasing dramatically the frequency of tumor-initiating cells. Mechanistically, MGP-driven OC stemness was mediated by the stimulation of Hedgehog signaling, in particular through the induction of the Hedgehog effector GLI1, thus highlighting a novel MGP/Hedgehog pathway axis in OCSC. Finally, MGP expression was found to correlate with poor prognosis in OC patients, and was increased in tumor tissue after chemotherapy, supporting the clinical relevance of our findings.

Thus, MGP is a novel driver in OCSC pathophysiology, with a major role in stemness and in tumor initiation.

## Introduction

Ovarian cancer (OC) is the deadliest gynecological tumor, with 313,959 new cases and 207,252 deaths worldwide in 2020 [1]. High-grade serous OC (HGSOC) is the most frequent type of OC (70%) with a survival rate of only 30% at 5 years [2]. Due to the lack of specific symptoms and screening methods, OC is frequently diagnosed at an advanced stage, when the tumor has already disseminated into the peritoneum. The standard of care for HGSOC is surgical cytoreduction followed by platinum-based chemotherapy. However, 70% of HGSOC relapse within 2 years [1], and almost all recurrent HGSOCs ultimately develop chemoresistance and become unresponsive to standard treatments [3]. Hence, new strategies to prevent, delay or treat tumor relapse in HGSOC patients remain an unmet need.

The clinical evolution of the disease seems to be accounted for by a small population of cells, called ovarian cancer stem cells (OCSC) [3]. Their stem-like features enable OCSC to initiate metastasis, drive tumor recurrence, and recapitulate the heterogeneity of the original tumor [4]. Furthermore, OCSC are able to elude the cytotoxicity of chemotherapeutics entering a quiescent state, increasing molecular pumps to efflux drugs outside the cells, and enhancing DNA repair and DNA damage response [5]. Hence, OCSC have emerged as optimal targets for novel OC-eradicating treatments. However, a detailed characterization of the molecular and functional traits of OCSC is still missing, partly due to the lack of clinically relevant experimental models. In fact, earlier studies on OCSC have mostly relied on cell lines that hardly recapitulate the biomolecular and histopathological features of HGSOC [3].

We have set up a pipeline that enabled us to explore the transcriptomes of patient-derived primary OCSC versus their non-stem counterpart. We found that one of the most prominent OCSC-associated genes was Matrix Gla Protein (MGP), which encodes an extracellular matrix protein that belongs to the vitamin K-dependent protein family. It is secreted by chondrocytes and vascular smooth muscle cells and is expressed in vessels, cartilage, bone, lung, heart and kidney [6]. MGP is classically known as a calcium chelator, mainly associated with the inhibition of tissue calcification in skeleton, coronary artery and kidney [7]. MGP has also been found aberrantly expressed in different cancer types [8], including OC [9], where its level of expression often correlates with tumor aggressiveness [10-12]. A few studies have also proposed a causal relationship between MGP expression and malignancy [13-15]. Yet, the biological mechanisms that regulate the functional contribution of MGP to cancer progression remain elusive. In particular, no information is available on the role of MGP in cancer stem cells (CSC), and also its involvement in OC development has not been investigated. Here, we show for the first time that MGP plays a pivotal role in OCSC pathophysiology. By altering the transcriptome of OCSC, MGP promoted stemness, epithelial-mesenchymal transition and tumor initiation. Furthermore, the use of patient-derived organotypic models revealed that MGP expression in OCSC is enhanced by the peritoneal microenvironment, thus pointing to this protein as a novel mediator of microenvironment-regulated OC stemness.

## Results

### MGP is upregulated in patient-derived OCSC

Primary cell cultures were established from tumor samples derived from HGSOC patients. The formation of single cell-derived spheroids under non-adherent conditions was employed in order to enrich for OCSC [16]. Patient-matched bulk, adherent tumor cells and OCSC-enriched spheroids were subjected to Affymetrix analysis and the transcriptomes were compared. A total of 2689 transcripts corresponding to 1176 genes (p<0.001, two-sample t-test with random variance model) were found to be differentially regulated between bulk and OCSC, with 560 genes upregulated and 616 genes downregulated in OCSC (Table S1). Among the upregulated genes we found MGP (FC=5.0, FDR=0.002; Fig. 1A), a gene that encodes for Matrix Gla Protein (MGP). Since MGP has never been associated with OCSC and very little is known about its function in cancer, we decided to investigate whether it plays any role in OCSC pathophysiology. First, we confirmed by RT-qPCR, in six patient-derived samples, the upregulation of MGP in OCSC as compared to their bulk counterpart (Fig. 1B). The data were also validated at the protein level, both by mass spectrometry-based proteomics in ten patient-derived samples (Fig. 1C and Suppl. Fig. S1A) and by western blotting in three patient-derived samples (Fig. 1D). Thus, MGP is consistently enriched in patient-derived OCSC. We also attempted to test whether MGP expression correlates with stemness-associated genes in HGSOC transcriptome datasets. To this goal, we performed a single-sample gene set enrichment analysis (ssGSEA) from 370 HGSOC patients included in the TCGA cohort. Overall, a positive and significant correlation could be found between MGP and gene sets related to stemness (Fig. 1E). Furthermore, the correlation extended also to gene sets related to epithelial-mesenchymal transition (EMT), which is a common feature of cancer stem cells [17]. In this regard, it is noteworthy that also in the context of the hallmarks’ gene sets MGP expression correlated with EMT (Fig. 1F).

**Figure 1.**
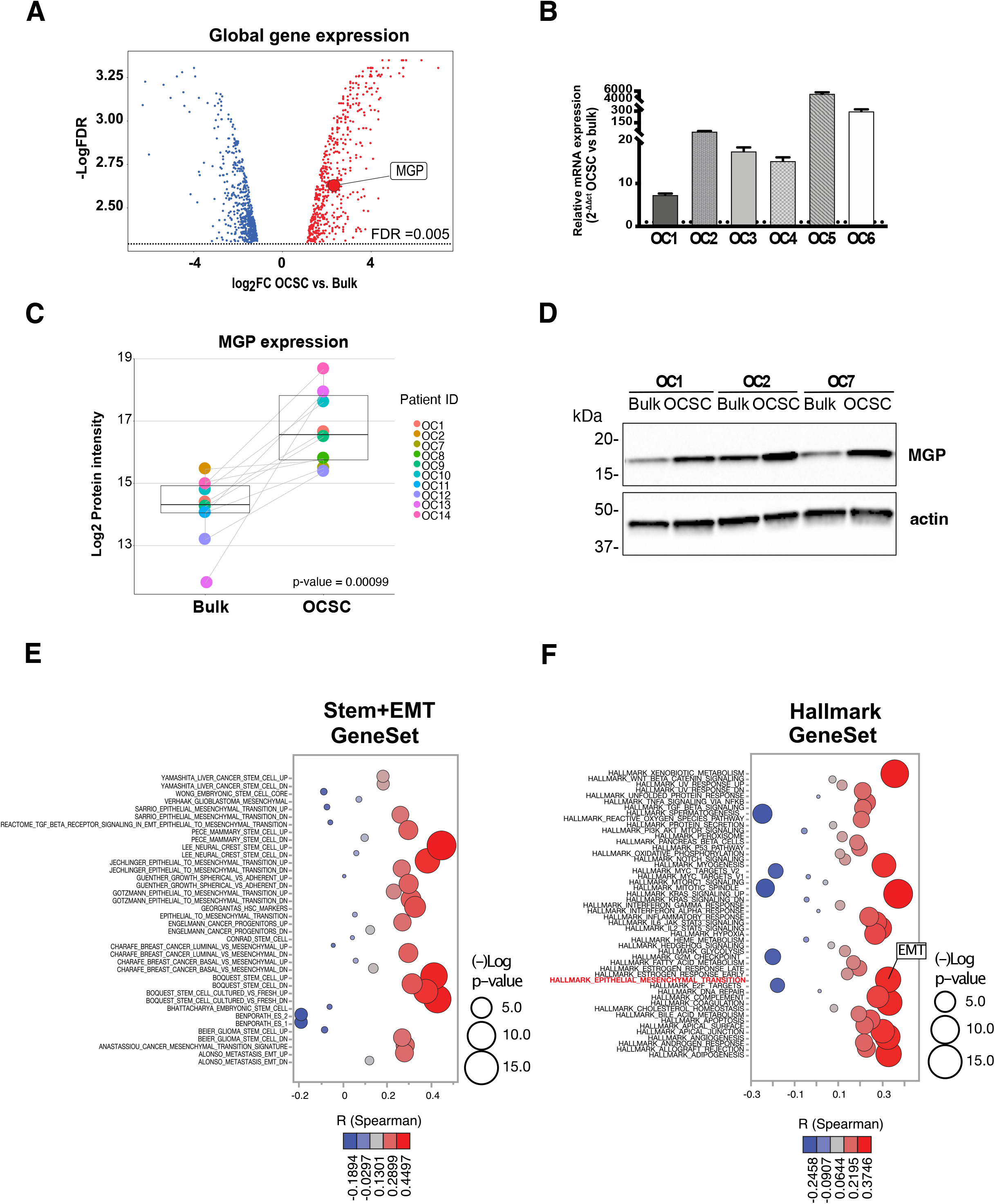
Upregulation of MGP in patient-derived OCSC. (A) Volcano plot showing the significantly differentially expressed genes (False Discovery Rate (FDR) ≤ 0.005) between primary spheres (OCSC) and adherent (bulk) cells. The x-axis represents the log2 fold change (log2FC; OCSC vs. bulk cells). DEGs with log2FC ≥1 and log2FC≤-1 are shown in red and blue, respectively. The y-axis represents the -Log False Discovery Rate (-Log FDR). MGP is indicated among the genes upregulated in OCSC. (B) MGP mRNA expression was analyzed by qRT-PCR. Data are represented as a relative mRNA expression (2^-Δ Δ Ct^) of MGP cultured as spheres as compared to their bulk counterpart (dashed line). (C) Normalized protein intensity for the protein MGP in ten patient-derived pairs of primary bulk OC cells and OCSC. The p-value was calculated by two-sided paired two-sample t-test. (D) Primary ovarian cancer cells from three independent patients (OC1, OC2 and OC7) were grown under bulk or OCSC conditions. Cell lysates were immunoblotted for MGP and actin was used as loading control. (E,F) Bubble plots showing ssGSEA results using the STEM+EMT GeneSets (panel E; N=38) or the HALLMARK GeneSets (panel F; N=50). Y-axes, GeneSets; X-axes, Spearman correlation coefficient (R) between MGP expression and ssGSEA scores in the TCGA-HGSOC cohort. Bubble size is proportional to inverse of Logarithmic (-Log) of significance [e.g., -Log(p-value) =5, p-value <10^−6^; student’s t-test] of Spearman correlation coefficient (R). Bubble colours refer to Spearman correlation coefficients (R), as per the legend.

### MGP is necessary and sufficient for OCSC features

To identify experimental models suitable to explore the functional role of MGP in OCSC, the protein level of MGP was evaluated in a panel of OC cell lines that have been classified as HGSOC-like models [18, 19]. We found that COV318, COV362 and Kuramochi cells expressed no or very low amounts of MGP, whereas the protein was readily detectable in Tyk-nu, OVCAR3 and OV90 cells (Suppl. Fig. S1B). Thus, COV318 and OVCAR3 were selected for the genetic manipulation of MGP in gain- and loss-of-function experiments, respectively. Endogenous MGP expression in OVCAR3 was ablated through the CRISPR-Cas9 technology and validated by RT-qPCR, western blot and immunofluorescence (Suppl. Fig. S1C-E). The ectopic expression of MGP in COV318 was achieved via lentiviral transduction, followed by validation (Suppl. Fig. S1F-H). Notably, neither knockout nor overexpression of MGP showed any effect on cell proliferation in standard 2D cultures (Suppl. Fig. S1I).

To investigate the contribution of MGP to OC stem-like features, we first took advantage of the sphere formation assay, which reflects the ability of CSC to resist to anoikis, to self-renew and to proliferate when cultured at low density under non-adherent conditions, generating monoclonal sphere-like structures [20]. The ablation of MGP expression significantly reduced the sphere-forming ability of OVCAR3, while MGP overexpression enhanced the spherogenicity of COV318 (Fig. 2A and Suppl. Fig. S2A). This demonstrated that MGP is causally involved in sphere formation and implicated for the first time MGP in OC stemness.

**Figure 2.**
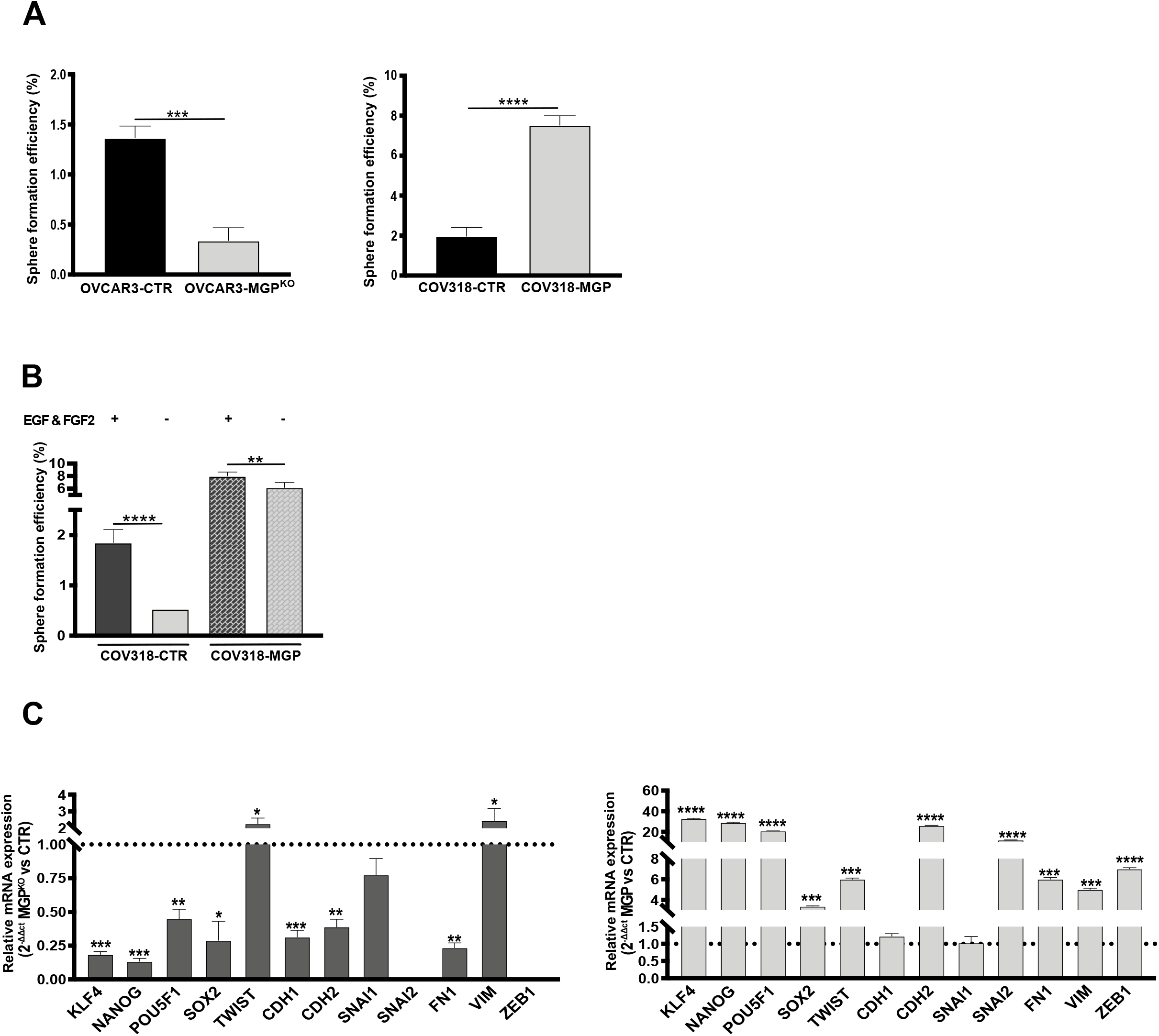
MGP modulates stemness-related features. (A) Sphere formation assay on second-generation spheres performed on MGP-manipulated cell lines. The experiment was conducted in triplicate and data are expressed as means ± SD. ***p<0.001, ****p<0.0001. (B) Sphere formation assay carried out on COV318-CTR or COV318-MGP in medium depleted of both EGF and FGF2. **p<0.01, ****p<0.0001. (C) qRT-PCR for stemness and EMT-related genes on second-generation spheres from OVCAR3 (left) and COV318 (right). Data are expressed as relative mRNA level (2^-ΔΔCt^) and normalized to the corresponding control cultures (dashed line). Data refer to means ± SD from a representative experiment performed in triplicate.

EGF and FGF2 are required for sphere formation in various stem cell models [16, 21, 22]. Surprisingly, we discovered that COV318-MGP cells, but not COV318-CTR, were able to generate spheres in the absence of growth factors (Fig. 2B), indicating that MGP is able to enhance OCSC function even under stringent conditions.

As a first attempt to elucidate the molecular mechanisms that underlie the role of MGP in the stem-like traits of OC cells, we tested whether MGP regulates the expression of genes involved in key aspects of stemness, namely multipotency and epithelial-mesenchymal transition (EMT). Indeed, the knock-out of MGP decreased the expression of several multipotency-associated genes, such as *KLF4, NANOG, POU5F1* and *SOX2*, while the ectopic expression of MGP induced their upregulation (Fig. 2C). A similar MGP-dependent regulation was observed for a subset of EMT-associated genes, including *CDH2, SNAI2, FN1* and *ZEB1* (Fig. 2C).

Taken together, these results support the notion that MGP confers stem-like traits to OC cells.

### MGP promotes OC cell adhesion and invasion of the mesothelium

Adhesion to and invasion of mesothelium are key steps in the peritoneal dissemination of OC, and OCSC are thought to play a pivotal role in this context, due to their ability to self-renew and to initiate the formation of metastatic and/or recurrent tumors [3]. Therefore, we evaluated whether MGP influences the adhesiveness of OCSC to the mesothelium. OCSC derived from second-generation spheres of OVCAR3 or COV318, with MGP knockout or ectopic expression, respectively, were seeded on top of a mesothelial layer formed by MeT5A cells, and the number of attached cells was measured after 8 hours. OVCAR3-MGP^KO^ cells exhibited a reduced adhesion ability compared to OVCAR3-CTR cells, while COV318-MGP showed significantly higher adhesion respect to COV318-mock (Fig. 3A). These data causally implicate MGP in the mesothelial adhesion of OCSC.

**Figure 3.**
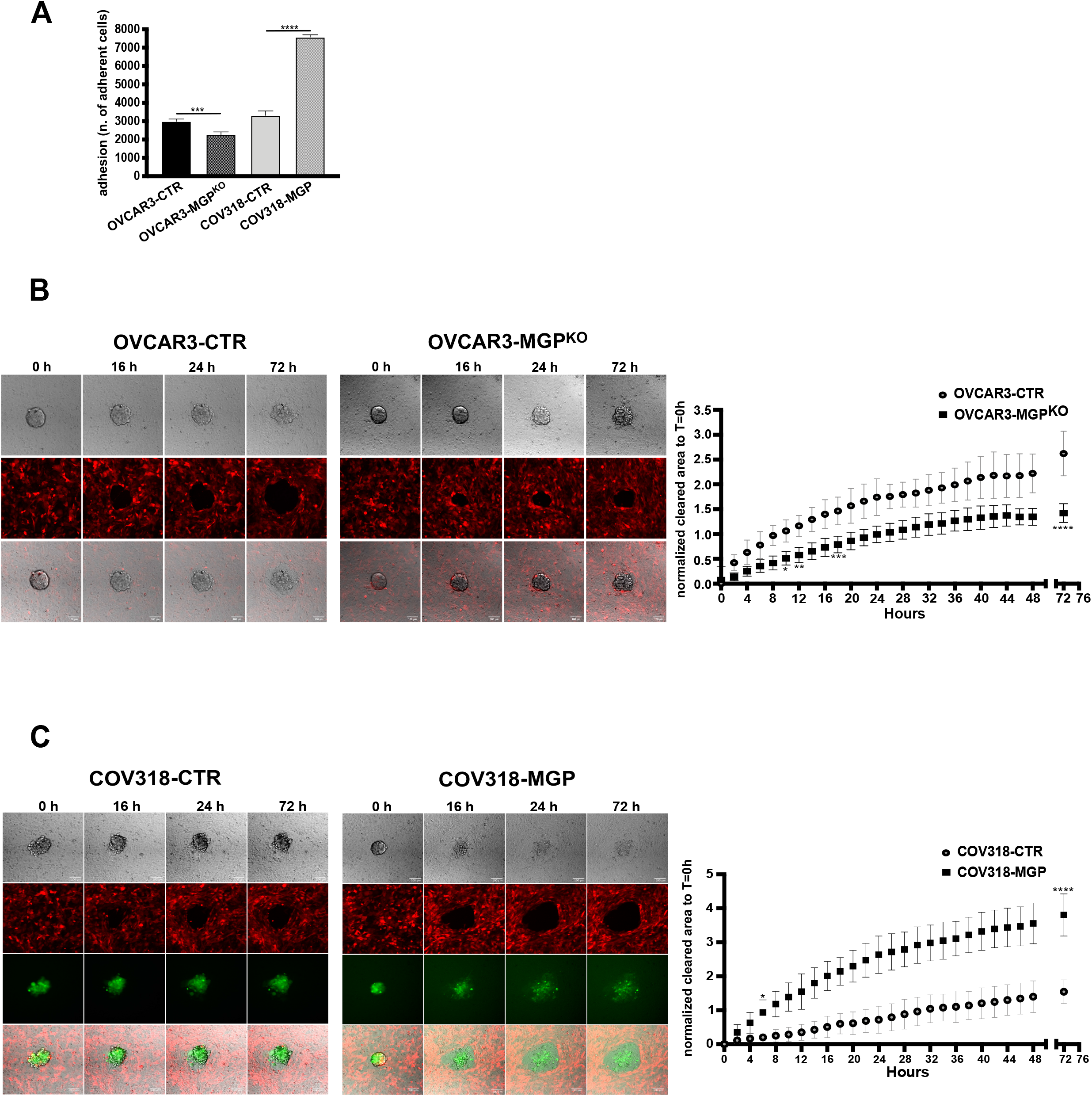
MGP regulates mesothelial invasion of ovarian cancer cells. (A) Adhesion of OVCAR3 (control vs MGP-knockout) or COV318 (control vs MGP-overexpressing) onto a monolayer of MeT5A-RFP cells. Results are expressed as number of adherent cancer cells as counted 8 hours after seeding. Data are expressed as means ± SD from two independent experiments each performed in sestuplicate. ***p<0.001 and ****p<0.0001. (B,C) Mesothelial clearance assays of OVCAR3 CTR or MGP^KO^ (B) and COV318 CTR or MGP (C). Images were taken at the indicated time points. The graphs on the side show the quantification of the cleared areas calculated every 2 hours (from 0 to 72 hours). Data are shown as means ± SD from 26 aggregates for each condition obtained from 3 independent experiments. * p<0.05, ** p<0.01, *** p<0.001, **** p< 0.0001. Scale bar, 100 µm.

To evaluate the functional role of MGP in mesothelial invasion, we employed the mesothelial clearance assay that recapitulates the early steps of OC peritoneal colonization [23]. Cell aggregates derived from OVCAR3-MGP^KO^ or COV318-MGP cells (or their respective controls) were plated on the mesothelial monolayer and their ability to breach and invade the monolayer itself was monitored over three days. As shown in Fig. 3B and in Suppl. videos 1 and 2, MGP ablation resulted in a decreased invasion of OVCAR3 cells, while the ectopic expression of MGP in COV318 cells enhanced their mesothelial invasion capability (Fig. 3C). The quantitation in Fig. 3B,C (right panels) shows that MGP exerted quite a dramatic effect on the mesothelial clearance potential of OC cells.

### MGP is upregulated by the TME and is required for TME-enhanced sphere formation

In an attempt to elucidate the biological basis of MGP enrichment in OCSC, we built on the notion that the tumor microenvironment (TME) influences several aspects of OC stemness [24] and, therefore, could be involved in the regulation of MGP expression in OCSC. We first tested this hypothesis in a clinically relevant setting consisting of an *in vitro* 3D organotypic model, derived from the patient’s omentum, that recapitulates the peritoneal TME [25]. Patient-derived primary OC cells were co-cultured with the 3D TME (Fig. 4A) followed by the RT-qPCR assay for the expression of MGP. In primary cultures from four independent patients, a remarkable upregulation of MGP was observed in cancer cells co-cultured with the TME as compared with cells that had no contact with the TME (Fig. 4B). Analogous effects were observed on a panel of OC-derived cell lines, including OVCAR3 and COV318 (Suppl. Fig. S2B). The peritoneal TME-dependent induction of MGP expression in OC cells implicates the TME in the regulation of cancer stemness. Along this line, the co-culture with the TME did indeed enhance the sphere-forming ability of OVCAR3 cells (Fig. 4C). Of note, MGP ablation abrogated sphere formation both in the absence and in the presence of TME (Figs. 2A and 4C), which implicates the molecule in both TME-induced and cell-autonomous spherogenesis.

**Figure 4.**
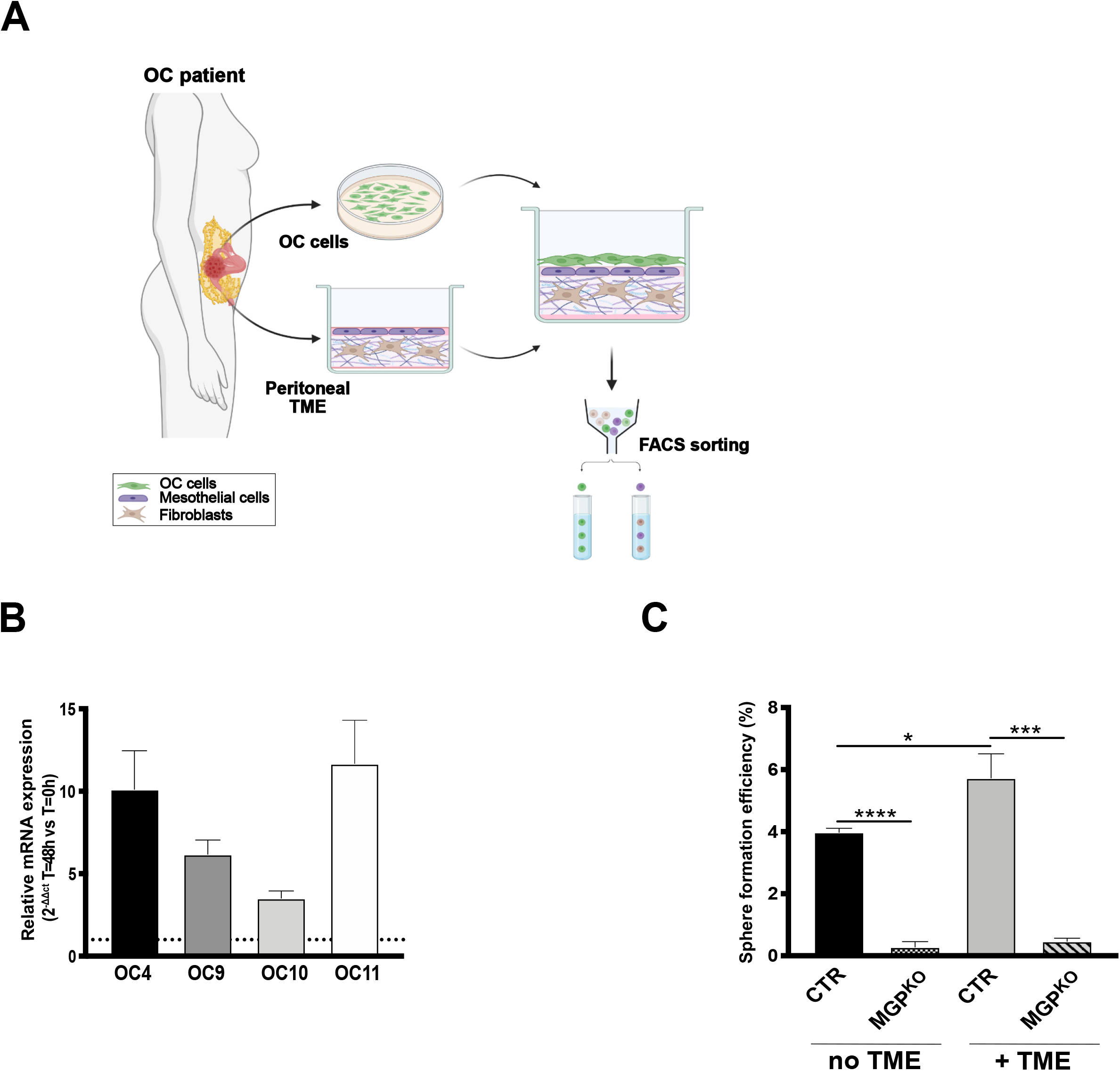
MGP in ovarian cancer cells is upregulated by the tumor microenvironment. (A) Schematic representation of the workflow used for the co-cultured of OC cells with the tumor microenvironment. (B) qRT-PCR for MGP on primary ovarian cancer cells from four independent patients co-cultured with the TME for 48 hours. Data are expressed as relative mRNA level (2^-ΔΔCt^) and normalized to the corresponding cells not co-cultured with TME (dashed line). Data refer to means ± SD from a representative experiment performed in triplicate. (C) Sphere formation assay on OVCAR3-CTR and OVCAR3-MGP^KO^ after a 48-hour co-culture with TME or without.

### MGP is involved in OCSC-driven tumorigenesis

Since tumor initiation is a defining feature of CSC, we asked whether MGP modulates the tumor-initiating potential of OC cells. NOD/SCID/IL2Rgamma-null mice were inoculated subcutaneously with OVCAR3 or COV318 genetically manipulated for MGP expression and monitored for tumor development.

We first determined the rate of tumor take in OVCAR3-CTR vs. OVCAR3-MGP^KO^ cells (5*10^6^ cells/mouse). As shown in the Fig. 5A, a dramatic reduction in tumor initiation was observed upon ablation of MGP: at 100 days post-injection, 10 out of 10 mice with OVCAR3-CTR cells developed tumors, while no tumors were observed in the MGP-knockout group. Furthermore, at day 200 only 2 mice out of 10 injected with OVCAR3-MGP^KO^ cells developed tumors.

**Figure 5.**
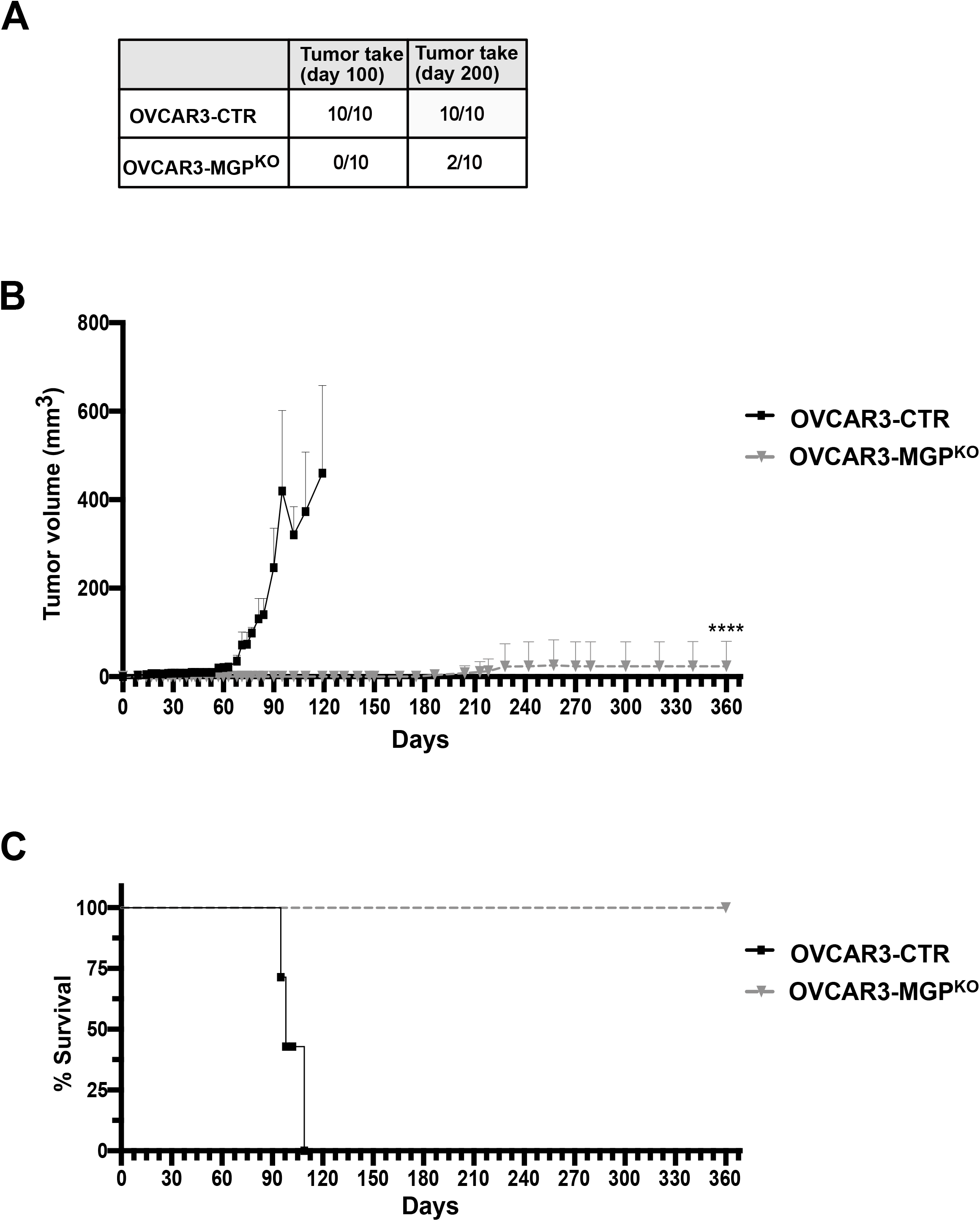
MGP is required for tumor initiation and growth. (A) OVCAR3-CTR or OVCAR3-MGP^KO^ were injected subcutaneously into NOD-SCID mice at 5*10^6^ cells/mouse (n=10). Tumor take was assessed at day 100 and 200 and is expressed as mice with palpable tumors. (B) Tumor growth was monitored at different time points. Data are expressed as mean tumor volumes ± SEM. **** p<0.0001. (C) Kaplan Meier survival analysis of mice injected with either OVCAR3-CTR or OVCAR3-MGP^KO^ cells. **** p<0.0001.

We observed a dramatic difference also in tumor growth, with control tumors reaching an exponential growth phase starting on day 60, whereas the tumors in the two mice with OVCAR3-MGP^KO^ cells failed to develop beyond a barely palpable size (Fig. 5B). Accordingly, all mice with OVCAR3-MGP^KO^ cells were all alive after 360 days, in contrast with the short survival of OVCAR3-CTR mice (Fig. 5C). Thus, MGP was required for both tumor initiation and growth.

These results raised the hypothesis that MGP increases the frequency of tumor-initiating cells in a population of OC cells. To test such a possibility, we performed *in vivo* extreme limiting dilution assays (ELDA) [26] with COV318 cells expressing MGP ectopically. Mice were injected with decreasing numbers of cells, ranging from 1*10^6^ to 10 cells/mouse, and then monitored for tumor formation. In agreement with the results on genetically manipulated OVCAR3 cells, MGP increased dramatically the tumor-initiating potential of COV318 cells. Indeed, tumor formation was observed at day 20 with as few as 500 cells in MGP-expressing cells, while 2*10^5^ cells were necessary to detect tumors with control COV318 cells at the same time point (Fig. 6A). ELDA calculations revealed that the frequency of tumor-initiating cells increased from 1/296,618 in COV318-CTR to 1/277 in COV318-MGP cells (1071 times; Fig. 6A), in line with MGP promoting a dramatic expansion of cells with tumorigenic capacity.

**Figure 6.**
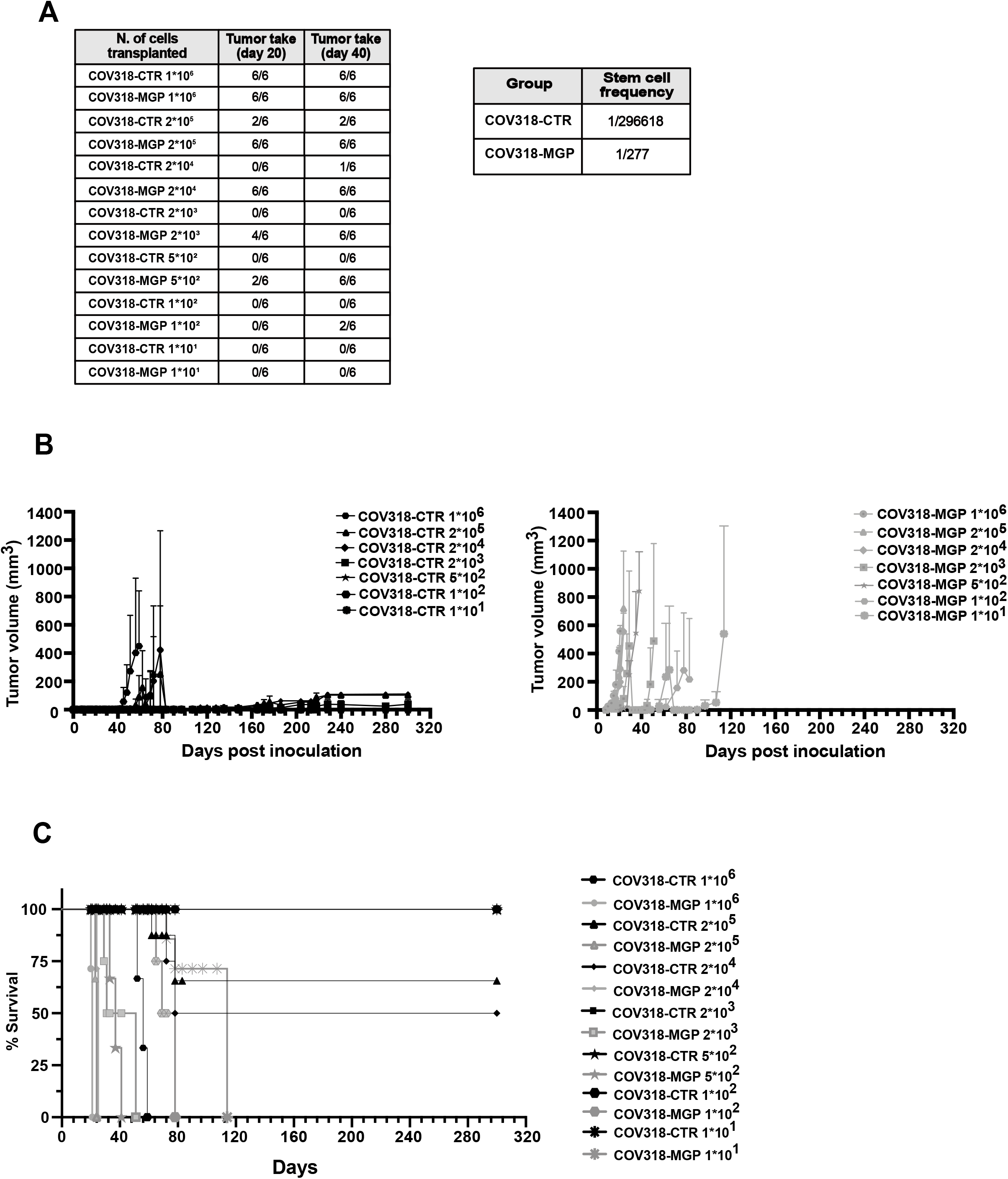
MGP accelerates tumor initiation, increases cancer stem cell frequency and enhances tumor growth. (A) NOD-SCID mice were transplanted subcutaneously with decreasing numbers of either COV318-CTR or COV318-MGP cells, and analyzed for tumor take at the indicated time points. Extreme limiting dilution assay (ELDA) was carried out to calculate the stem cell frequency (p=5.81*10^−31^). (B) The same mice were monitored over time and tumor volumes were measured. Data are expressed as means ± SEM. **** p<0.0001 (C) Kaplan–Meier survival analysis of mice injected with decreasing numbers of either COV318-CTR or COV318-MGP cells. **** p<0.0001, *** p<0.001 for 10 cells.

Besides tumor initiation, the ectopic expression of MGP also enhanced and sustained COV318 tumor growth. Among COV318-CTR mice, only the animals injected with 1*10^6^ and a fraction of those injected with 2*10^5^ or 2*10^4^ cells developed tumors, while lower cell numbers formed either no tumors or palpable lesions that remained small even after 10 months (Fig. 6B, left). On the contrary, in the COV318-MGP group all mice developed tumors, including those inoculated with 10 cells, and all tumors exhibited exponential growth (Fig. 6B, right). Accordingly, COV318-MGP tumors contained a remarkably higher frequency of proliferating, Ki67-positive cells as compared to control tumors (Suppl. Fig S2C). Along the same line, COV318-MGP tumors resulted in short survival even with low cancer cell numbers, while survival was markedly longer for COV318-CTR mice, with animals being still alive at 10 months when injected with up to 2*10^4^ cells (Fig. 6C).

These data indicated a major role for MGP in tumor initiation and development. Taken together, both gain and loss-of-function approaches pointed to MGP as a driver in OCSC pathophysiology and revealed that MGP is not only associated to the OCSC phenotype but is actually implicated in OCSC-driven malignancy.

### MGP promotes OC stemness via GLI1

To elucidate the molecular mechanisms underlying the role of MGP in OCSC, and to identify downstream effectors, we first defined the impact of manipulating MGP expression on the transcriptome of these cells. RNA-seq analysis was carried out in MGP-manipulated vs. control cells, comparing bulk with OCSC. We found 1624 genes differentially regulated in bulk cells (Suppl. Fig. S2D and Table S3) and 999 in OCSC (Fig. 7A and Table S4) that showed opposite trend of regulation upon either MGP ablation in OVCAR3 cells or overexpression in COV318 cells, thus implicating MGP as a master modulator of OC transcriptome. We then applied a reverse engineering approach based on EnrichR to identify molecular mechanisms modulated by MGP. This revealed several tumor-associated pathways in both bulk (Suppl. Fig. 2E) and in OCSC (Fig. 7B), with the latter enriched in pathways classically implicated in stemness (*e*.*g*., EMT, WNT/β-catenin, estrogen response). A subset of these pathways was also identified by a GSEA approach (Suppl. Fig. S2F).

**Figure 7.**
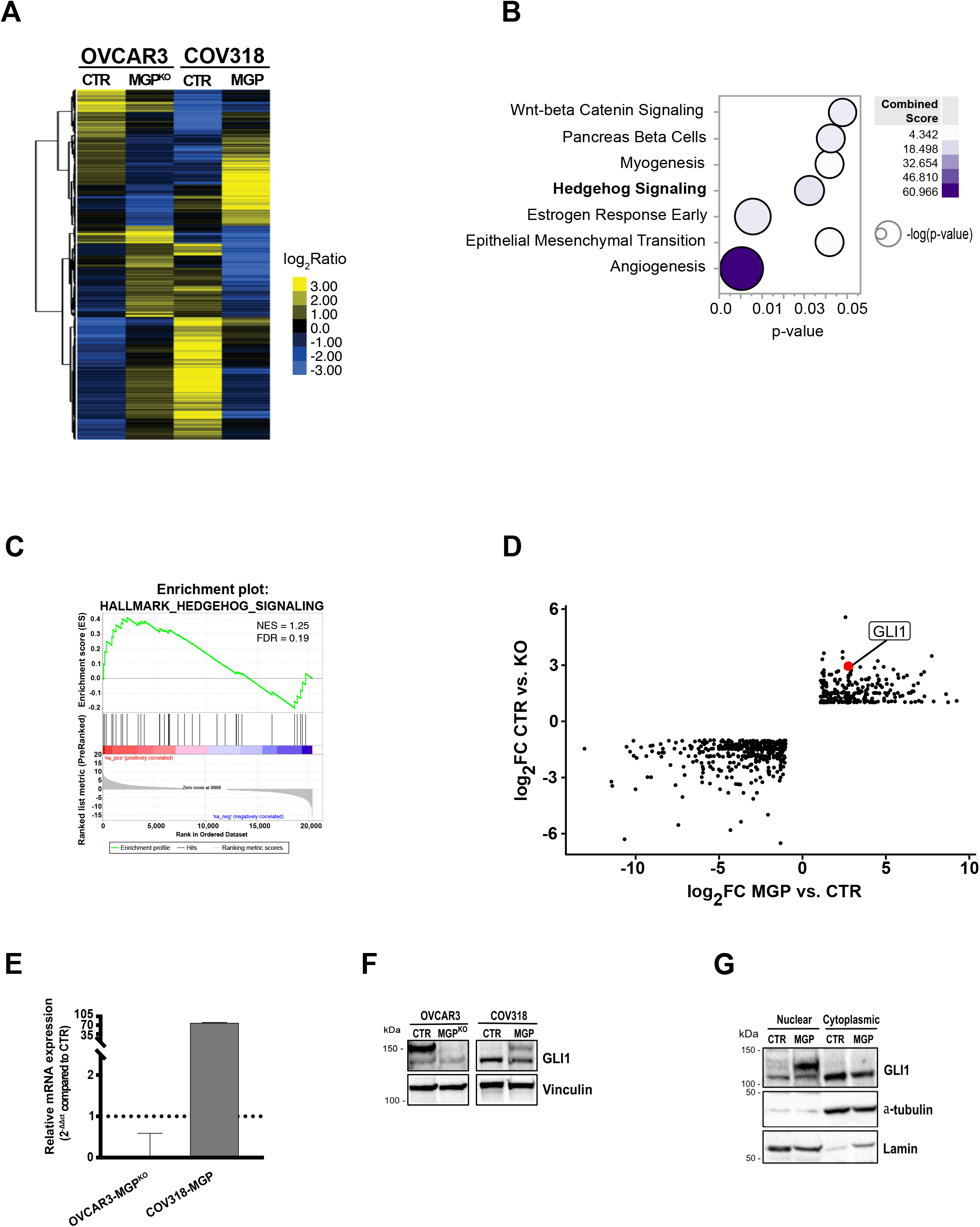
MGP regulates gene expression in OCSC. (A) Hierarchical clustering analysis of MGP regulated genes in OVCAR3-CTR vs. OVCAR3-MGP^KO^ and COV318-CTR vs. COV318-MGP cells cultured as OCSC. The heatmap shows the expression of differentially regulated genes. The relative log_2_FC ratio values of expression are indicated in the legend. (B) Bubble plot showing the EnrichR analysis using the HALLMARK GeneSets (N=38) in COV318-MGP OCSC with opposite trend compared to OVCAR3-MGP^KO^. Y-axes, GeneSets; X-axes, p-value of enrichment significance. Bubble size is proportional to inverse of Logarithmic (-Log) of p-value. Bubble colours refer to the combined score value, as per the legend. (C) Pre-ranked GSEA enrichment plots showing hallmark of cancer related to Hedgehog signaling pathway in COV318 cells cultured as OCSC. NES, normalized enrichment scores; FDR, significance of the enrichment (False Discovery Rate; 1000 random permutations); p, nominal p-value; ‘na_pos’ and ‘na_neg’ phenotypes correspond to enrichment in upregulated and downregulated genes, respectively. (D) Scatter plot showing the differentially expressed genes between OCSC overexpressing or knock-out for MGP. The x-axis represents the log2 fold change (log2FC) of OVCAR3-MGP^KO^ vs. OVCAR3-CTR cells. The y-axis represents the log2FC of COV318-CTR vs. COV318-MGP cells. The red dot indicates the position of GLI1. (E) qRT-PCR for GLI1 on OVCAR3-CTR vs. OVCAR3-MGP^KO^ and COV318-CTR vs. COV318-MGP cells. Data are expressed as the relative mRNA expression (2^-ΔΔCt^) and normalized to the corresponding control cells (dashed line). Data refer to the means ± SD from a representative experiment performed in triplicate. (F) Immunoblot for GLI1 on MGP-manipulated OVCAR3 and COV318 cells. Vinculin was used as loading control. (G) Immunoblot for GLI1 on cytoplasmic and nuclear fractions from COV318-CTR vs. COV318-MGP cells. Alpha-tubulin and Lamin were used as loading controls for cytoplasmic and nuclear fractions, respectively.

One of the most prominent pathways modulated by MGP in OCSC was Hedgehog signaling (Fig. 7B and Suppl. Fig. S2F), a pathway that has been implicated in stem-like properties of OC cells [27] but for which a link with MGP has never been reported. A GSEA confirmed the MGP-induced activation of the Hedgehog pathway in OCSC (Fig. 7C). Accordingly, the transcription factor GLI1, a major effector of Hedgehog signaling, was among the top-ranking genes regulated by MGP in OCSC (Fig. 7D). The upregulation of GLI1 in MGP-overexpressing cells and its downregulation in MGP-knockdown cells was validated at both the mRNA and the protein levels (Fig. 7E, F). Since the activation of GLI1 upon Hedgehog pathway stimulation is a direct consequence of its nuclear translocation, we tested the intracellular localization of the protein through subcellular fractionation. As shown in Fig. 7G, the overexpression of MGP resulted in massive accumulation of GLI1 in the nucleus of COV318 cells, further supporting the connection between MGP and Hedgehog signaling. We then asked whether GLI1-mediated Hedgehog signaling is involved in the regulation of OC stemness by MGP. The activation of the Hedgehog pathway can occur in a canonical, Patched1/Smoothened-mediated manner (PTCH1-SMO-GLI1; [28]) or in a non-canonical way, which refers to the SMO-independent activation of GLI1 [29]. GANT61, a specific GLI1 inhibitor which targets both canonical and non-canonical Hedgehog pathway [30], reduced MGP-driven, but not basal, sphere formation in a dose-dependent manner (Fig. 8A). Analogous results were obtained with cyclopamine (Fig. 8B), which influences the balance between the active and inactive forms of Smoothened and inhibits specifically the canonical pathway [31]. Notably, the cyclopamine derivative vismodegib, an FDA-approved Hedgehog inhibitor [32], also reduced MGP-dependent sphere formation dose-dependently (Suppl. Fig. S2G). Thus, MGP-induced sphere formation in OC cells occurred through the canonical Hedgehog pathway. To further elucidate the role of Hedgehog signaling as an effector of the stem-like phenotype promoted by MGP, we tested whether GLI1 is involved in the induction of multipotency and EMT-related genes. As shown in Fig. 8C, GANT61 efficiently reduced the MGP-dependent upregulation of *KLF4, NANOG, POUF1* and *SOX2*, as well as of several genes associated with EMT (*TWIST1, CDH2, SNAI2, FN1, VIM, ZEB1*). On the contrary, the same genes were not affected by the GLI1 inhibitor in control cells.

**Figure 8.**
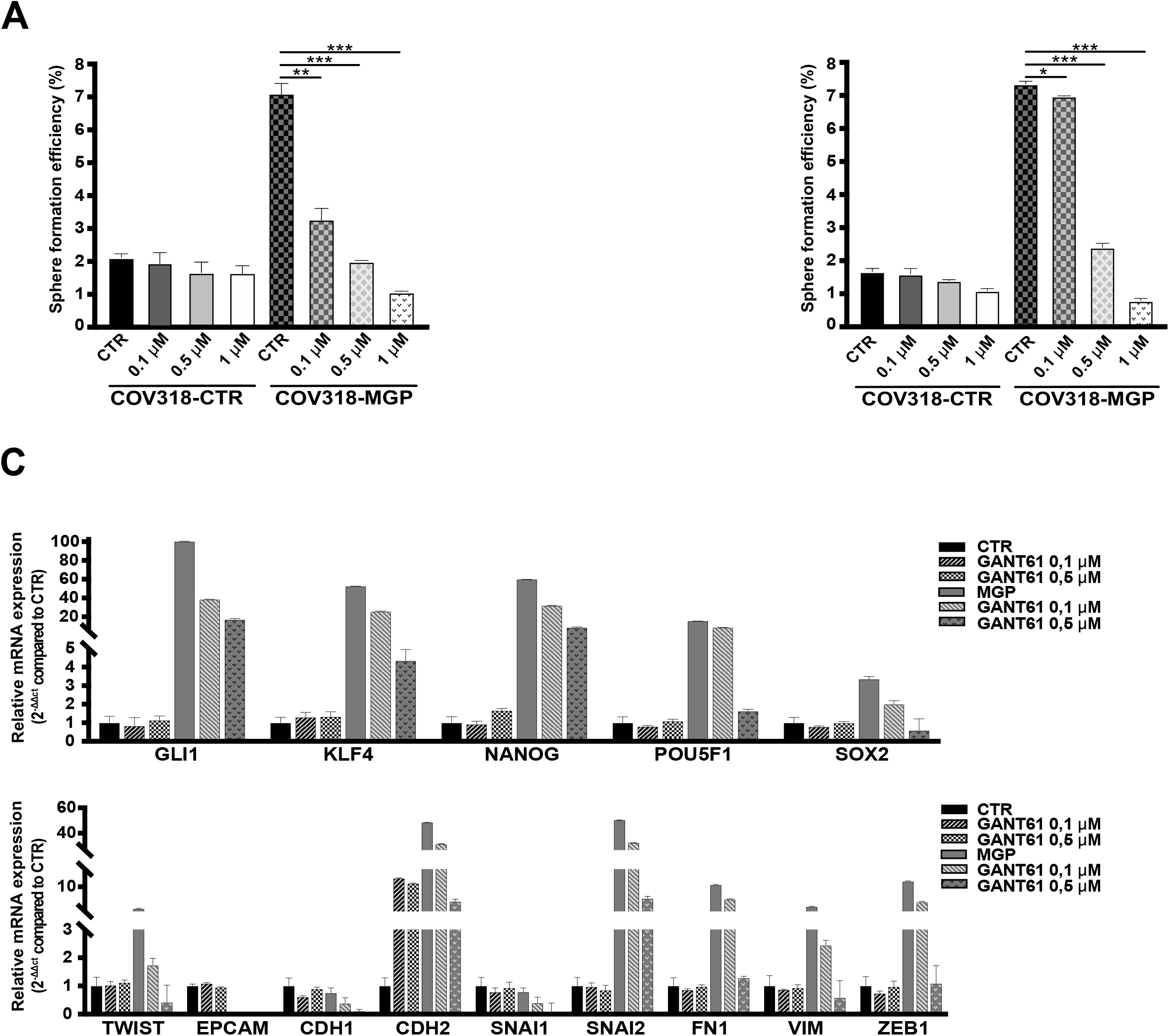
MGP-induced stemness is mediated by Hedgehog signaling. (A,B) COV318-CTR and COV318-MGP cells were treated with different concentrations of GANT61 (A) or cyclopamine (B) and subjected to sphere formation assay. For each analysis, results are shown as means ± SEM from two independent experiments. ** p<0.01, *** p<0.001. (C) qRT-PCR for stemness (top graph) and EMT-related genes (bottom graph) on COV318-CTR and COV318-MGP treated with 0.1 or 0.5 µM GANT61 for 48 hours. Data are expressed as relative mRNA level (2^-ΔΔCt^) and normalized to the COV318 control culture. Data refer to means ± SD from a representative experiment performed in triplicate.

Taken together, these results implicate GLI1-mediated Hedgehog signaling in the stem-like phenotype of OC induced by MGP.

### The clinical relevance of MGP in OC

Our experimental data pointed to MGP as a driver of aggressiveness in OC. To clarify to what extent this applied to the human disease, we first analyzed the expression of the protein in tissue samples from HGSOC or from the fimbriated end of healthy fallopian tubes, the main tissue of origin for this tumor type [33]. Notably, MGP was detected at variable levels in all tumor samples. In contrast, no or low expression of MGP was found in healthy fimbriae (Fig. 9A). We then investigated the prognostic implications of MGP expression in OC by testing the correlation between MGP levels and the survival time in HGSOC patients from the TCGA cohort. Indeed, high MGP correlated significantly with shorter overall survival (HR 1.72; Fig. 9B, left). A similar trend was observed for disease-free survival, although statistical significance was not reached (HR 1.21; Fig. 9B, right). These data revealed that MGP is associated with poor prognosis in HGSOC patients. Of note, the *MGP* gene maps on 12p12.3, a hotspot for copy number alterations in OC (data available at https://www.cbioportal.org/) and, in fact, the gene was amplified in 8% of ovarian cancer in the TCGA cohort (Suppl. Fig. S3A). In agreement with the gene expression data reported in Fig. 9B, patients with *MGP* gene amplification had shorter overall survival than patients with no amplification (Suppl. Fig. S3B).

**Figure 9.**
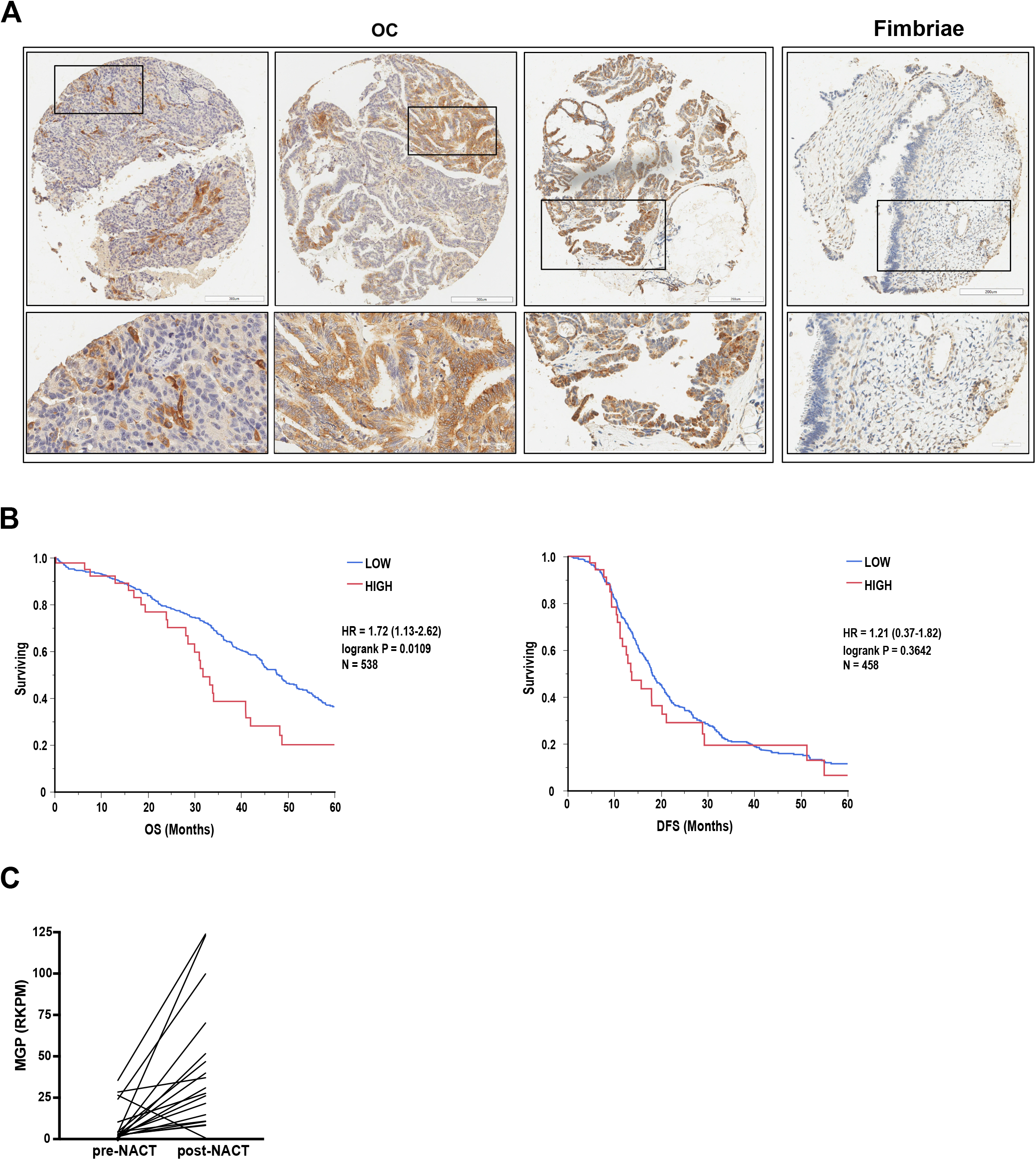
Clinical relevance of MGP in human OC. (A) Sections of 3 representative OC samples and 1 fimbriae sample stained for MGP. Sections were counterstained with Hematoxylin. Lower panels show higher magnifications of the insets indicated in upper panels. Scale bars: 300 µm, upper panels; 50 µm, lower panels. (B) Survival analysis of TCGA-HGSOC patients stratified by MGP gene expression level (see methods). HIGH = high expression of MGP; LOW = low expression of MGP. Panels show overall survival (OS) on the left and disease-free survival (DFS) on the right. Log-rank p-values are displayed together with hazard ratio (HR, 95% Confidence Interval) and number of patients. (C) Expression of MGP in 19 patients before and after neoadjuvant chemotherapy (NACT). The data are obtained from the RNA-sequencing dataset published by Javellana et al [37], and are expressed as reads per kilobase million (RPKM). *** p<0.001.

Finally, given that chemotherapy has been proposed to promote stem-like traits in OC cells [34-36], we investigated the expression levels of MGP in paired samples of OC collected from 19 patients before and after chemotherapy [37]. MGP was found to be consistently upregulated in post-treatment samples (Fig. 9C), suggesting that chemotherapy promotes an enrichment in MGP-expressing tumor cells.

## Discussion

While MGP alterations have been conclusively implicated in pathological calcification processes, their association with, and functional contribution to, neoplastic diseases remain controversial. For example, the downregulation of MGP has been proposed as a hallmark of colon carcinoma [38], but other investigators reported that colon cancer progression and worse prognosis are rather associated with higher MGP expression [14]. Contradictory results have also been published in glioblastoma, where both the upregulation and downregulation of MGP have been linked to tumor aggressiveness [39]. In general, whether MGP correlates with higher or lower aggressiveness depends on the tumor type or subtype. This is best exemplified by breast cancer, where MGP is downregulated in advanced HER2-positive tumors [40] while high levels are associated with the aggressive triple-negative subtype [13]. With regard to ovarian cancer (OC), Hough et al. reported that MGP is upregulated in this tumor type, yet with no indication on its prognostic value. We now confirmed and extended that observation by showing that the protein is detectable at variable level in OC but not in the tissue of origin, and its expression correlates with shorter survival in OC patients. The clinical value of MGP is further supported by our finding that MGP levels in OC are consistently increased upon treatment with chemotherapy, in agreement with the notion that the protein is associated with a highly malignant phenotype.

Our results strongly suggest that MGP upregulation is not a mere epiphenomenon of OC aggressiveness, but it rather reflects a functional contribution to tumor development. In particular, we report a novel role of MGP in the pathophysiology of OCSC and propose that such a role underlies the pro-tumorigenic function of MGP. By analogy to many other solid tumors, also in OC the sub-population of CSC have been implicated in various aspects of disease progression. This paradigm seems particularly applicable to the clinical evolution of OC, which is characterized by high rate of metastatic spread, high frequency of relapse and development of chemoresistance, all features commonly ascribed to CSC. Accordingly, various studies have identified neoplastic cells with stem-like traits that fuel OC spread and recurrence (reviewed in [3]). Nevertheless, the molecular and functional properties of OCSC have remained largely elusive due to different factors, one of the most prominent being the lack of clinically representative model systems. We have overcome this issue by relying on a series of patient-derived primary cultures, an approach that has recently revealed novel and unexpected players not only in OC *per se* but also in the OCSC subset [16, 41-43]. Along this line, primary cultures unveiled MGP as a hallmark of OCSC and provided the rationale for subsequent functional studies.

We could therefore show for the first time that MGP acts as a driver of cancer stemness, a function that entails the orchestration of a set of genes classically involved not only in dedifferentiation and multipotency but also in EMT, which is considered as an intrinsic feature and a causal player in the CSC phenotype [17]. Based on these findings, it is conceivable that the functional contribution of MGP to cancer stemness is not limited to OC. While, as outlined above, the role of MGP in cancer aggressiveness remains controversial and tumor type-dependent, we argue that in those cancers where MGP appears to exert a pro-tumorigenic function, such a function could entail the activation of the CSC subpopulation. Along the same line, future studies should test whether MGP expression represents a proxy for the “stemness” degree of a given tumor. This could have important therapeutic implications, especially in view of the emerging treatments that are being developed against CSC [44].

One of the most striking and unexpected findings in our study is the strong regulatory function exerted by MGP on the transcriptional activity of OC, with over 1600 genes showing opposite regulation upon MGP ablation vs. ectopic expression in bulk tumor cells and almost 1000 in OCSC. Such a massive regulation of the transcriptome, which is ascribed to MGP for the first time, most likely accounts for the pleiotropic function of the protein that we found in OC stemness. At the same time, the modulation of so many pathways that are known to play pivotal roles in tumor progression (e.g., hypoxia, angiogenesis, STAT5, etc.; Figs. 7B and S2E) raise the hypothesis that the role of MGP in OC extends well beyond stemness and orchestrates various aspects of cancer development. For example, the regulation of STAT5- or angiogenesis-related pathways might reflect the involvement of MGP in the crosstalk of tumor cells with the immune or the vascular systems, respectively. In agreement with the possible involvement of MGP in various cancer-related processes not necessarily related to OCSC, we observed a widespread distribution of the protein (i.e., not restricted to small cancer cell subpopulations) in various OC specimens within our tissue microarrays (Fig. 9A).

Our study identified the Hedgehog pathway as a prominent effector of MGP-driven OC stemness, and implicated Hedgehog signaling in particular in self-renewal, and in the expression of multipotency and EMT-related genes. These findings are consistent with the reported tumor-promoting effect of Hedgehog activation in OC [45]. Notwithstanding, an early-phase clinical trial in OC patients with the Hedgehog inhibitor vismodegib has not given satisfactory results [46]. While this failure may depend on several factors, our observations raise the intriguing possibility that tumors with higher MGP levels and, hence, more sustained Hedgehog pathway activation, are more responsive to Hedgehog inhibitors. Therefore, patient stratification according to MGP levels may help identifying subsets of individuals who are more likely to benefit from the treatment with Hedgehog pathway inhibitors.

MGP has been classically described as a secreted protein with extracellular activity, especially in the context of preventing ectopic calcification of the extracellular matrix [7]. However, we consistently failed to detect MGP in the extracellular environment of OC cells, including both the soluble compartment and the extracellular matrix (not shown). This implies that the multiple processes regulated by MGP in OCSC rely on intracellular activities of the protein. The only study published so far which reported an intracellular function of MGP was conducted on gastric cancer cells [15]. Interestingly, those investigators implicated intracellular MGP in the activation of STAT5 signaling, which is one of the pathways we found enriched in MGP-expressing OC cells (Fig. 1F). While these observations might point to the stimulation of the STAT5 pathway as a general hallmark of intracellular MGP activity in cancer cells, it should be presumed that the molecular routes that underlie the MGP/STAT5 crosstalk are different. In fact, while Wang et al. reported STAT5 activation to be the consequence of the nuclear translocation of MGP [15], we only detected MGP in the cytosol of OC cells with no evidence of nuclear localization (Suppl. Figs. S1E,H and data not shown). Although beyond the scope of the present study, future work should focus on how cytosolic MGP exerts its pleiotropic role in OC, and particularly in OCSC.

In this context, we demonstrated for the first time that MGP induces GLI1-mediated Hedgehog signaling, a pathway that has been implicated in various aspect of OC progression including cancer stemness. Aberrant Hedgehog signaling, indeed, is associated with reduced recurrence-free or overall survival in OC patients [47, 48]. The ectopic activation of the pathway in OC cells, for instance by GLI1 overexpression, promotes cell proliferation, cell mobility and invasiveness [48]. GLI1 activity has also been reported to confer stem-like traits to OC cells, a function that is mediated by the multipotency-associated transcription factor NANOG [49]. Of note, NANOG is dramatically upregulated by MGP in our OCSC models thus raising the possibility of a MGP-GLI1-NANOG axis as a driver of OC stemness. Our finding that MGP-induced upregulation of NANOG is partially reverted by a GLI1 inhibitor (Fig. 8C) would support this hypothesis. Our platform of fully patient-derived organotypic models revealed that the expression of MGP in OC cells is induced by the contact with the peritoneal TME, that represents the most prominent site for metastasis and recurrence in this tumor type. We also provided evidence that the TME enhances sphere formation in OC cells. These findings are consistent with the emerging view that a crosstalk with the peritoneal niche is essential for several stemness-associated features of OC, such as tumor initiation, metastasis, relapse and therapy resistance [24]. Our data, besides providing further support to such a paradigm, suggest that regulating MGP expression is an integral part of the OCSC-orchestrating function of the peritoneal TME. It is intriguing that MGP, in turn, promotes the peritoneal adhesion and invasion of tumor cells, which points to this protein as a master regulator of the bi-directional crosstalk between the TME and OCSC. On a more general level, defining the TME-derived factors that induce and/or sustain OCSC activity in the context of metastasis and tumor recurrence will be instrumental to identify new actionable targets for OC eradication.

## Conclusion

We have unveiled the novel and unexpected role of MGP as a master regulator of the pathophysiology of OCSC, as mirrored not only by its ability to promote self-renewal, the expression of stemness genes, and massive transcriptome changes, but also by a remarkable tumor-initiating potential. While providing new mechanistic insights into the pathogenic function of OCSC, our findings pave the way to further investigation on MGP and its downstream effectors in this context, which, in turn, may help designing innovative therapeutic strategies to defeat OCSC-driven tumor progression.

## Declarations

### Ethical Approval

Informed consent was obtained from the patients enrolled in the study. This study was performed in accordance with the Declaration of Helsinki and was approved by the European Institute of Oncology Ethics Committee (protocol no. R789-IEO). Animal studies were performed following protocols approved by the fully authorized animal facility of European Institute of Oncology and by the Italian Ministry of Health (as required by the Italian Law) (protocol no. 325/2016) and in accordance with EU directive 2010/63.

### Authors’ contributions

V.N., J.V.O, F.B. and U.C. contributed to the concept and design of the study. V.N., C.B., G.F, M.L. and C.S. performed experiments. N.C. provided clinical specimens. V.M, G.F., G.B and F.B. contributed to the analysis and interpretation of results. V.N. F.B. and U.C. wrote the manuscript. All authors reviewed and approved the manuscript.

### Availability of data and materials

All materials area available from the authors upon reasonable request. Affymetrix raw and normalized data are available in Gene Expression Omnibus (GEO) with the accession number GSE216627 (https://www.ncbi.nlm.nih.gov/geo/query/acc.cgi?acc=GSE216627). RNA sequencing data are available in Gene Expression Omnibus (GEO) with the accession number GSE216487 (https://www.ncbi.nlm.nih.gov/geo/query/acc.cgi?acc=GSE216487). Raw mass spectrometry data are available at http://proteomecentral.proteomexchange.org via the PRIDE partner repository with the dataset identifier PXD037543.

## Abbreviations

CSC: Cancer stem cells
ELDA: Extreme limiting dilution assay
EMT: Epithelial-mesenchymal transition
FC: Fold change
FDR: False discovery rate
GSEA: Gene set enrichment analysis
HGSOC: High-grade serous
OC MGP: Matrix Gla Protein
OC: Ovarian cancer
OCSC: Ovarian cancer stem cells
TCGA: The cancer genome atlas
TME: Tumor microenvironment

## Materials and Methods

### Cell culture

Human ovarian carcinoma cell lines OVCAR3 were purchased from American Type Culture Collection (ATCC; cat# HTB-161), COV318 were acquired from Sigma (cat# 07071903). OVCAR3 cells were grown in RPMI 1640 medium (Euroclone, cat# ECM2001L) containing 20% fetal bovine serum (FBS, cat# RM10432), 2 mM L-glutamine (Lonza, cat# 17-605E), 100 U/ml penicillin, 100 μg/ml streptomycin (Euroclone, cat# LODE17602E), 10 µg/ml bovine insulin (Merck, cat# 91077C). COV318 cells were cultured in DMEM (Euroclone, cat# ECM0103L) supplemented with 10% FBS, 100 μg/ml streptomycin and 100 U/mL penicillin. The immortalized human mesothelial cell line MeT5A was kindly provided by Prof. Aldieri (Turin) and maintained in Medium 199 (Sigma, cat# M4530) containing 10% FBS, 3.3 nM epidermal growth factor (Merck, cat# E4127), 400 nM hydrocortisone (Sigma, cat# H0888), 870 nM Bovine insulin and 20 mM HEPES (Sigma, cat# H0887). Cell lines were routinely tested for mycoplasma with a PCR-based method and authenticated via short tandem repeat profiling.

Primary cell cultures were established from peritoneal ascites of high-grade serous ovarian cancer (HGSOC) patients. Samples were provided by the Division of Gynecology at the European Institute of Oncology (Milan) upon informed consent from patients undergoing surgery. Tumor histology was confirmed by a board-certified pathologist (GB), while the identity of cancer cells was confirmed by immunostaining for cytokeratins 5, 7, and 8, or for pan-cytokeratins. The purity of primary cell culture was consistently over 95%. Tissue isolation and culture conditions of primary cells were performed as described previously [41].

Mesothelial and fibroblasts were isolated from healthy omental specimens. The omentum was washed several times with sterile PBS, centrifuged at 1500 rpm for 5 minutes and cultured with RPMI 1640, 20% FBS and 100 U/ml penicillin, 100 μg/ml streptomycin. To isolate fibroblasts, the tissue was minced with scissors and razor and incubated overnight on an orbital shaker with 100 U of hyaluronidase (Merck, cat# H3884) and 1,000 U of collagenase type III (Worthington Biochemical, cat# LS004183) in DMEM, 10% FBS at 37°C. The tissue was then centrifuged at 1500 rpm for 5 minutes, washed two times with PBS and plated with DMEM, 10% FBS, 1% glutamine, 100 U/ml penicillin, 100 μg/ml streptomycin, 1% NEAA (Lonza, cat# 13-114E), 1% MEM vitamins (Lonza, cat# 13607C). Mesothelial and fibroblast cells were used for the experiments at early passages (1-3). All cell lines and primary samples were maintained at 37°C in a humidified incubator with 5% CO_2_.

### Sphere Formation Assay

Sphere formation assays were performed as described [16]. Briefly, single cell suspension derived from ovarian cancer cell lines or primary samples, were seeded at low density under non-adherent conditions in poly-(2-hydroxyethyl methacrylate) coated dishes (Sigma, cat# P3932-25G) and allowed to form monoclonal spheres. For ovarian cancer cell lines, OCSC-enriched sphere were maintained in DMEM:F-12 (1:1) (Gibco, cat# 11320033), supplemented with 2% B27 (Thermo Fisher Scientific, cat# 17504044;), 2 mM L-glutamine, 100 U/ml penicillin, 100 μg/ml streptomycin, 20 ng/mL EGF (Merck, cat# E4127), and 10 ng/mL fibroblast growth factor-2 (FGF2; Peprotech, cat# AF-100-18B). Cells were cultured at a density of 1000 cells/ml. Spheres were then dissociated with StemPro™ Accutase™ (Thermo Fisher Scientific, cat# A1110501), according to the manufacturer’s protocol, and re-plated under the same conditions to obtain second-generation spheres.

Primary OCSC cultures were seeded at 5000 cells/ml in MEBM™ (Lonza, cat# CC-3151) supplemented with 2 mM L-glutamine, 100 U/mL penicillin, 100 µg/mL streptomycin, 5 µg/mL insulin, 0.5 µg/mL hydrocortisone, 1 U/mL heparin (Voden, cat # 07980), 2% B27, 20 ng/mL EGF, and 20 ng/mL FGF2.

Sphere formation was assessed 5–10 days after seeding. The sphere-forming efficiency (SFE) was defined as the ratio between the number of spheres counted and the number of cells seeded.

### Affymetrix Analysis

The Affymetrix analysis was performed as described [16] on patient-matched adherent cells and second-generation spheres derived from the 7 patients described in Table S2. After RNA extraction, samples were grouped into two independent pools of 4 and 3 samples, respectively. Total RNA was isolated using the miRNeasy Micro kit (Qiagen, cat# 217084) according to manufacturer’s protocol and quantified using a NanoDrop 2000 (Thermo Scientific). The Ovation® Pico WTA System V2 (Nugen, cat# 3302-60) was used to amplify 5 ng of total RNA according to manufacturer’s protocol. Quality control of the RNA samples was performed using an Agilent Bioanalyzer 2100 (Agilent Technologies). Each pool was labeled and hybridized to Affymetrix® Human Gene 2.1 ST according to the manufacturer instructions (Affymetrix). Quality control and normalization of Affymetrix.CEL files using the Robust Multi-array Average (RMA) were performed by Expression Console software (Affymetrix; version: 1.4.1.46). Microarray data analysis was performed using the BRB-ArrayTools (Version 4.4.0). Gene expression data were log2 transformed and probesets (psets) whose variance was in the bottom 75^th^ percentile were excluded, for a total of 13404 psets retained in the analysis. Class comparison analysis to identify differentially regulated genes was performed using t-test with a random variance model. The number of psets significant at 0.001 level of the univariate test was 2689 for a total of 1134 unique genes.

### RNA extraction, qRT-PCR and sequencing

Total RNA was extracted from adherent cultures or from spheres using the RNeasy Mini Kit (QIAGEN, cat# 217004) according to the manufacturer’s protocol and quantified using the 2100 Bioanalyzer (Agilent). RNA quality control was checked using an Agilent 2100 Bioanalyzer (Agilent, Santa Clara, CA, USA). Preparation and hybridization of cDNA samples were performed at the Cogentech Microarray Unit (Milan, Italy; www.cogentech.it).

Gene expression levels for MGP, EMT and stem genes were analyzed and normalized against housekeeping human GAPDH and HPRT1. TaqMan assays for specific genes are listed in Table S5. Normalized expression changes were determined with the comparative threshold cycle (2^−ΔΔCT^) method.

TruSeq kit and Illumina HiSeq2000 were used for RNA sequencing following the manufacturer’s instructions. The sequencing coverage and quality statistics for each sample are summarized in Table S6.

### RNA-Seq analysis

Poly-A enriched strand-specific libraries were generated with the TruSeq mRNA V2 sample preparation kit (Illumina, cat# RS-122-2001), ribosomal RNA depleted strand-specific RNA libraries with the TruSeq Stranded Total RNA LT sample preparation kit with Ribo-Zero Gold (Illumina, cat# RS-122-2301 and #RS-122-2302), and transcriptome capture based libraries with the TruSeq RNA Access Library Prep Kit (Illumina, cat# RS-301-2001). Recommended amounts of starting material were as follows: 100 ng of input RNA for TruSeq, 100 ng for Ribo-Zero, and 10 ng of intact RNA or 20 ng of degraded RNA for RNA access. All protocols were performed following the manufacturer’s instructions. Libraries were sequenced by Illumina HiSeq2000 resulting in paired 50nt reads. Fastq files were aligned to the hg38 genome assembly using STAR. STAR gene counts were normalized applying the median of ratios method implemented in DESeq2 R package. Briefly, the normalization process implies different steps: i) for each gene, a pseudo-reference sample is created and is equal to the geometric mean across all samples; ii) for every gene in a sample and for each sample, the ratios sample/ref are calculated; iii) the median value of all ratios for a given sample is taken as the normalization factor (size factor) for that sample; iv) for each gene in each sample the normalized count values is calculated dividing each raw count value by the sample’s normalization factor. For each gene, in each experimental condition (spheres cells and adherent cells), log2 Fold Change (log2FC) was calculated as the log2 ratio of normalized reads in MGP-overexpressing cells/KO cells vs. control cells.

Heatmaps and clusters were generated by using Cluster 3.0 for Mac OS X (C Clustering Library 1.56) and Java TreeView version 1.1.6r4 (uncentered correlation and centroid linkage) using median centered log2FC data. Volcano plot were generated using R and log2FC data. Pre-Ranked Gene Set Enrichment Analysis (https://www.gsea-msigdb.org/gsea/index.jsp) was performed using log2FC of COV318 spheres and adherent cells, 1000 random gene sets permutation, weighted class metric, and hallmark of cancer and C2 curated Molecular Signatures Database gene sets29,30 (MSigDB). EnrichR analysis was performed on upregulated genes on COV318-MGP cells which showed an opposite regulation compared to OVCAR3-MGP^KO^. We followed the instruction on the webtool https://amp.pharm.mssm.edu/Enrichr.

### Immunoblotting

Cells, both bulk and spheres, were lysed in lysis buffer (4% SDS, 16% glycerol, 40 mM Tris-HCl [pH 6.8]) for 15 minutes at 90°C, centrifuged for 5 minutes at 14,000 g and the supernatant was collected.

To prepare cytoplasmic extracts, the cells were washed with cold PBS and incubated with a buffer (10 mM Hepes pH 7.9; 1 M KCl; 0.1 mM EDTA; 0.1 mM EGTA, 1:500 protease and phosphatase inhibitors) for 20 minutes on ice in a shaker. The cells were then scraped and collected in an eppendorf tube. After centrifugation for 10 minutes at maximum speed, the supernatant containing the cytoplasmic fraction was collected. To prepare nuclear extracts, the pellet was washed 3 times with the same buffer and then incubated with the hypotonic buffer (20 mM Hepes pH 7,9; 400 mM NaCl; 0.1 mM EDTA; 0.1 mM EGTA, 50% Glycerol; 1:500 protease and phosphatase inhibitors) for 1 hour in a shaker at 4°C. Nuclei were collected by centrifugation for 5 minutes at maximum speed.

Protein concentration was determined using a Pierce BCA Protein Assay kit (Thermo Fisher Scientific, Inc, cat# 23227) according to the manufacturer’s instructions. Equal amounts of protein extracts (20 µg) were resolved in acrylamide gel and transferred onto nitrocellulose membranes. The membranes were incubated overnight at 4°C with the following primary antibodies: MGP (Abcam, cat# ab86233, 1:500), GLI1 (cell signaling, cat# 2643S, 1:1000), vinculin (Sigma-Aldrich, cat# V9131, 1:10,000), actin (Abcam, cat# ab853, 1:3000), GAPDH (Sigma, cat# ABS16, 1:2000), Lamin A/C (Santa Cruz, cat# sc-7292, 1:500), α-tubulin (Santa Cruz, cat# sc-32293, 1:1000). Membranes were incubated with IgG HRP-conjugated secondary antibody (Bio-Rad Laboratories, cat# 1706515, 1706516, dilution 1:3000) for 1 hour at room temperature. The signal was detected by the Clarity Western ECL Substrate (Bio-Rad, cat# 1705062) as described in the manufacturers protocol and the images were acquired using ChemiDoc (Bio-Rad) and analyzed with the Fiji software.

### Proteome analysis

Cells were lysed with boiling guanidine-hydrochloride lysis buffer (6 M Gnd-HCl, 100 mM Tris-HCl pH 8.5, 5 TCEP, 10 mM CAA). Lysates were heated at 95 °C for 10 minutes, while shaking, and sonicated (Vibra-Cell VCX130, Sonics, Newtown, CT) for 2 minutes, with pulses 1 seconds on, 1 second off. Protein concentrations were determined using the Bradford assay (Bio-Rad). Proteins were pre-digested with endoproteinase Lys-C (Wako) for 2 hours at room temperature in an enzyme/protein ratio of 1:100. Thereafter, samples were diluted to 1 M guanidine-hydrochloride with 25 mM Tris buffer before overnight digestion with trypsin (Sigma-Aldrich) at 37°C in an enzyme/protein ratio of 1:50. After overnight digestion, enzymatic activity was quenched by acidifying the lysates using trifluoroacetic acid (TFA) at a final concentration of 1% and ensuring the pH of the samples being around 2. 750 ng of digested peptides for single-shot proteome analysis were loaded on C18 Evotips (Evosep) for MS analysis. Two technical replicates per sample were prepared.

Samples were analyzed on the Evosep One system [50] coupled to an Orbitrap Exploris 480 [51]. Samples were separated on an in-house packed 15 cm analytical column (150 μm inner diameter), packed with 1.9 μm C18 beads, and column temperature was maintained at 60 °C using an integrated column oven (PRSO-V1, Sonation GmbH) on the 30 samples per day gradient. The mass spectrometer was operated in positive ion mode, with spray voltage at 2 kV, heated capillary temperature at 275 °C and funnel RF frequency at 40. Peptide match was set to off, and isotope exclusion was on. We used data-independent acquisition (DIA), with full MS resolution set to 120,000 at m/z 200 and full MS AGC target set at 300%, with an injection time of 45 ms. Scan range was set to 350– 1400 m/z. AGC target value for fragment scan was set at 1000%. 49 windows of 13.7 Da were used with an overlap of 1 Da. The MS/MS acquisition was set to 15,000 resolution, and injection time to 22 ms. Normalized collision energy was set at 27%.

DIA raw files were analyzed on Spectronaut [52] v 15.5.211111.50606 in directDIA mode with standard settings. Deamidation of asparagine and glutamine (NQ) was added as variable modification. Data filtering was set on “q value sparse” (default in version 15). The Human Uniprot fasta file (downloaded in October 2020; 20,600 entries) was supplemented with the MaxQuant contaminant fasta file containing 246 entries.

The plot shown in Fig. 1C was generated using the R software v4.1.1 (with RStudio v1.2.504) with the ggplot2 package v3.3.5.

### Immunofluorescence

OVCAR3 and COV318 cells were seeded on coverslips and grown to a near-confluent state. Cells were fixed with 4% paraformaldehyde for 10 minutes at room temperature and then permeabilized in ice-cold PBS, 0.5% Triton X-100 for 3 minutes at 4°C. After blocking for 1 hour at room temperature in a humid chamber with blocking buffer (PBS, 2% BSA, 5% donkey serum, and 0.05% Triton X-100), the cells were incubated for 2 hours with the anti-MGP primary antibody (Abcam, cat# ab86233, 1:50) diluted in blocking buffer. Coverslips were then washed with PBS and incubated with the secondary antibodies for 1 hour at room temperature (Jackson Immuno Research Laboratories, Cy5 Donkey anti-rabbit, cat# 711-175-152, 1:400). Nuclei were then counterstained with 0.2 µg/ml DAPI (Sigma-Aldrich, cat# 32670-25MG-F) and the coverslips were mounted in Mowiol (Merck, cat# 81381). Images were acquired using the Leica MultiFluo microscope.

### Cell viability

Ovarian cancer cells were seeded in 96-well plates (1.5*10^3^ cells/well in quadruplicate) and incubated for 24-48-72-96 hours. After each time point, the metabolic activity was quantified using the Cell counting kit-8 (Sigma-Aldrich, cat# 96992), following the manufacturer’s instructions. The absorbance at a wavelength of 450 nm was measured using the Glomax Plate Reader.

### Adhesion assay

MeT5A-RFP were seeded at 2*10^4^ cells/well into a 96 well plate and incubated for 24 hours at 37°C in order to obtain a cell monolayer. 7*10^3^ single cells derived from second-generation spheres were labeled with 1 mM CMFDA (Life Technologies, cat# C7025) for 30 minutes and added on top of the mesothelial monolayer. The cells were washed after 8 hours. The attached cells were imaged using a Nikon Eclipse microscope and counted using the Image J software.

### Mesothelial Clearance Assay

MeT5A-RFP were plated into a 96 well plate at 2*10^4^ cells/well and incubated for 24 hours at 37°C in order to obtain a cell monolayer. Ovarian cancer spheroids were formed by incubating 3*10^2^ cells/well in a poly-HEMA coated 96-well round-bottom plate at 37°C for 24 hours. The spheroids were then transferred onto the mesothelial monolayer and the co-cultures were imaged every 2 hours for 72 hours using a Nikon Eclipse microscope.

The invasive area created by the aggregates in the mesothelial monolayer was measured and the clearance was calculated normalizing the cell-free area created by the spheroid with the aggregate size at time 0. Representative images were selected and combined in Image J software to get the clearance videos.

### 3D organotypic model

The 3D organotypic model was assembled by plating 6*10^4^ primary omental fibroblasts mixed with 7.5 µg of collagen-I and 7.5 µg of fibronectin at 37°C. After 4 hours, 3*10^5^ primary mesothelial cells were added to the culture and incubated at 37C for 2 days. Single-cell suspensions of primary HGSOC cells, pre-labeled with 1 µM CMFDA for 15 minutes, were seeded on top of 3D organotypic cultures. After 48 hours, the co-cultures were dissociated to single cells and CMFDA-labeled HGSOC cells were isolated by FACS sorting and subjected to RNA extraction.

When the 3D organotypic model was established using cell lines, IMR90 (fibroblasts), MeT5A (mesothelial) and OVCAR3 (either control or MGP^KO^) were employed, and after FACS sorting tumor cells were subjected both to RNA extraction and to sphere formation assays.

### Cell treatments with Hedgehog inhibitors

Second-generation spheres from COV318-CTR or COV318-MGP cells were seeded at 1*10^3^ cells/ml in triplicate in polyHEMA-coated 6-well plates. Sphere culture and SFE measurement were performed as described above.

The following Hedgehog inhibitors were used in sphere formation assays at the concentrations indicated in the figures: GANT61 (Selleckem, cat# S8075), Cyclopamine (Fisher scientific, cat# J61528.MB), and Vismodegib (Selleckem, cat# S1082). SFE was determined 7 days after treatment.

### Immunohistochemistry

Immunohistochemical (IHC) staining from human tissue microarray or mouse xenograft samples was performed on 3-μm-thick sections from formalin-fixed paraffin-embedded samples and dried in a 37°C oven overnight. Hematoxylin/eosin (H&E) staining was performed with H&E Leica Kit Infinity for a full automated autostainer (Leica ST5020). IHC staining for Ki67 and MGP antibodies was performed using Bond III IHC autostainer for full Automated Immunohistochemistry (Leica biosystems). Antigen was unmasked with Tris-EDTA pH 9 (Bond Epitop Retrival Solution 2 Leica, cat# AR9640). Both mouse monoclonal anti-Ki67 CloneMIB-1 (Dako, cat# M7240) and rabbit polyclonal antibody MGP (ProteinTech, cat# 10734-1-AP) were used at 1:200. The antibodies were diluted with Bond Primary Antibody Diluent AR9352 Leica. BOND IHC Polymer Detection Kit (cat# DS9800) was used to stain both antibody with DAB Cromogen. IHC samples were counterstained using Hematoxylin solution (Leica, cat# RE7107-CE). Pictures of stained sections were acquired with the Aperio ScanScope XT instrument. Ki67 and MGP staining were analyzed and scored by a board-certified pathologist (GB).

### Lentivirus production and cell transduction

For lentiviral infection, HEK293T was used as the packaging cell line. HEK293T were co-transfected, using the calcium phosphate precipitation method, with 10 µg of lentiviral vectors either empty vector (EX-NEG-Lv122; GeneCopoeia) or containing the cDNA for human MGP (EX-Z9471-Lv122; GeneCopoeia), and the following packaging vectors: PMD2G, RRE and REV.

The supernatant from HEK293T containing the virus particles was supplemented with 8 µg/mL of polybrene and used to infect COV318 cells, generating the COV318-CTR and COV318-MGP cell lines upon selection with 2.5 µg/ml puromycin. Overexpression of MGP was confirmed by Western blot analysis and immunofluorescence.

### CRISPR-Cas9

Single guide RNA (sgRNA) sequences to target MGP gene in the exon 1 region were designed using the online web tool for CRISPR (https://chopchop.cbu.uib.no; [53]). Two sgRNA were selected (GCCTTCCACTAACATCCCGTAGG and CAAAGTTACTACCGCTAAGGCGG) in order to perform a dual guide targeting approach. Both guide sequences were then cloned as DNA inserts into pSpCas9 (BB)-2A-GFP (pX458) (Addgene plasmid, cat# 48138; a gift from Bruno Amati, Milan, Italy), encoding also the Cas9 protein and GFP.

To generate stable OVCAR3 knock-out cell lines, 10 µg pX458 encoding Cas9 and sgRNAs were transfected using Lipofectamine 3000 (ThermoFisher, cat# L3000008), according to manufacturer’s instruction. Sorting of GFP-positive cells was performed 48 hours after transfection and 1*10^3^ cells were plated onto a 15-cm dish. Clones were isolated 10-15 days later and MGP-knockout clones were identified by PCR and Sanger sequencing.

### *In vivo* models

6-8 week-old NOD/SCID IL2R-gamma null (NSG) female mice (from Charles River Laboratories) were injected subcutaneously into the flank with ovarian cancer cells in a 1:1 (vol:vol) mixture with growth factor-reduced Matrigel (Corning, cat# 356231) and phosphate-buffered saline (PBS), with a final volume of 100 µl per injection. Each experimental group consists of 6 mice for COV318 or 10 mice for OVCAR3.

For limiting dilution experiments, mice were injected with serial dilutions of COV318-CTR or COV318-MGP cells ranging between 1*10^6^ and 10 cells/site, while 5*10^6^ OVCAR3-CTR or OVCAR3-MGP^KO^ cells/site were injected into the mice flank.

Tumor latency was defined as the time interval from the injection to the formation of palpable tumors. Tumor take was determined as number of mice with palpable tumors. Tumor size was monitored by caliper measurement and the growth curves of different tumors were calculated with the formula: Tumor Volume = 1/2*(length × width^2^).

Body weight and general physical status were monitored daily, and the mice were sacrificed when the tumor reached 1.2 cm in diameter. Survival curves were drawn using the Kaplan-Meier method. The log-rank Mantel-Cox test was employed to define any statistical difference between the survival curves of the groups. The stem cells frequency was measured using the ELDA online Software http://bioinf.wehi.edu.au/software/elda.

All experimental procedures involving mice and their care were conducted in conformity to the following laws, regulations and policies governing the care and use of laboratory animals: Italian Law (D.lgs 26/2014, authorization no. 19/2008-A issued 6 March 2008 by the Ministry of Health); internal protocol approved by the fully authorized animal facility of European Institute of Oncology and by the Italian Ministry of Health (IACUC no. 25/2015). Mice were housed under specific pathogen-free conditions in isolated vented cages and allowed access to food and water ad libitum.

### TCGA-HGSOC data analysis

Total RNA-seq gene expression data (N=307) and clinical data for TCGA patients with high-grade serous ovarian adenocarcinoma (TCGA-HGSOC) available at cBIOportal (https://www.cbioportal.org/) were used. For single-sample gene set enrichment analysis (ssGSEA) of RNA-seq TCGA cohort, enrichment scores were calculated by using the GSVA package in [54] and gene sets of Molecular Signatures Database (MSigDB) [55, 56] related to stemness (STEM), epithelial-to-mesenchymal transition (EMT) and hallmarks of cancer. Spearman correlation coefficients between ssGSEA results and MGP expression, and p-values were calculated using R (R Core Team (2021). R: A language and environment for statistical computing. R Foundation for Statistical Computing, Vienna, Austria. URL https://www.R-project.org/). For survival analyses we downloaded MGP copy number variants classification (CNV, N=579) and MGP microarray gene expression data z-score classification (z-score cut-off for high expression = +1.5, N=538) for TCGA patients with high-grade serous ovarian adenocarcinoma (TCGA-HGSOC) available at cBIOportal (https://www.cbioportal.org/). Overall survival (OS) and disease free survival (DFS) were estimated by the Kaplan-Meier method. Follow-up was truncated at 5 years in order to reduce the potential overestimation of overall mortality with respect to ovarian-cancer specific mortality. All survival analyses were performed using the JMP software, version 16 (SAS Institute Inc., Cary, NC, 1989–2021).

## Data Availability Statement

Affymetrix and RNAseq data were deposited in Gene Expression Omnibus database (http://www.ncbi.nlm.nih.gov/geo/) with the accession numbers GSE216627 and GSE216487, respectively.

Raw mass spectrometry data have been deposited to the ProteomeXchange Consortium (http://proteomecentral.proteomexchange.org) [57] via the PRIDE partner repository [58] with the dataset identifier PXD037543.

## Acknowledgements

We are grateful to all patients who donated their samples for this work. We thank A. Gatto for technical assistance with primary cultures, G. Jodice and S. Pece for the immunohistochemical analyses, E. Lengyel and H. Kenny for kindly providing omental primary cells. P. Nicoli for assistance with the mouse experiments. We thank the staff of Division of Gynecology and of Biobank at IEO for providing clinical specimens.

Fabrizio Bianchi wishes to dedicate this work to the memory of Lucia, an extraordinary girl who experienced cancer as an opportunity to entrust her and all her loved ones to the love of the Father.

