## Supplementary Figures and Legends for "Matrix Gla Protein acts as a driver of stemness and tumor initiation in ovarian cancer"

#### Supplementary figure legends

##### Supplementary Figure 1

(A) Extract ion chromatogram (XIC) relative to the peptide YAMVYGYNAAYNR from MGP in patient OC9. iRT = indexed Retention Time. The chromatogram was generated in Spectronaut. (B) Immunoblot for MGP on a panel of ovarian cancer cell lines. Actin was used as a loading control. (C,D,E) Endogenous MGP expression in OVCAR3 cells was ablated through the CRISPR-Cas9 technology. Knockout of MGP was assessed by qRT-PCR (C), immunoblotting with vinculin as loading control (D), and immunofluorescence (E), with MGP in red and nuclei counterstained with DAPI (blue). (F,G,H) Ectopic expression of MGP in COV318 cell line was achieved via lentiviral transduction and was confirmed by qRT-PCR (F), immunoblotting with GAPDH as loading control (G), and immunofluorescence (H), with MGP in red and nuclei counterstained with DAPI (blue). (I) Growth curves of MGP-manipulated OVCAR3 and COV318 cells. Proliferation was evaluated by CCK-8 assay and the data show the means ± SD from 3 independent experiments. No significant difference was observed in either cell line upon MGP manipulation.

##### Supplementary Figure 2

(A) Sphere formation assay performed on second-generation spheres on two clones of OVCAR3-CTR and five clones of OVCAR3-MGP$^{KO}$. Data refer to a representative experiment performed in triplicate and are expressed as means ± SD. Comparisons between controls and knockout were done with two-sided Student's t-tests. ** $p<0.01$, *** $p<0.001$, **** $p<0.0001$. (B) qRT-PCR for MGP on a panel of ovarian cancer cell lines co-cultured with the TME for 48 hours. Data are expressed as relative mRNA level ($2^{-\Delta\Delta Ct}$) and normalized to the matched cell line that had no contact with TME (dashed line). The experiments were performed in triplicate with GAPDH and HPRT1 as normalizers. (C) IHC staining for MGP and Ki67 on tumor sections derived from mice injected with either COV318-CTR or COV318-MGP. Scale bar = 50 µm. (D) Hierarchical clustering analysis of MGP regulated genes in OVCAR3-CTR vs. OVCAR3-MGP$^{KO}$ and COV318-CTR vs. COV318-MGP cells cultured as bulk populations. The heatmap shows the expression of differentially regulated genes. The relative $log_2FC$ ratio values of expression are indicated in the legend. (E) Bubble plot showing the EnrichR analysis using the HALLMARK GeneSets (N=38) in COV318-MGP bulk cells with opposite trend compared to OVCAR3-MGP$^{KO}$. Y-axes, GeneSets; X-axes, p-value of enrichment significance. Bubble size is proportional to proportional to inverse of Logarithmic (-Log) of p-value. Bubble colours refer to the combined score value, as per the legend. (F) Bubble plot showing GSEA results using the HALLMARK GeneSets (N=50) in COV318-MGP OCSC cells. Y-axes, GeneSets; X-axes, nominal p-value of enrichment significance. Bubble size is proportional to inverse of Logarithmic (-Log) of false-discovery rate (FDR) of enrichment significance (1000 random data permutations). Bubble colours refer to normalized enrichment score (NES), as per the legend. (G) COV318-CTR and COV318-MGP cells were treated with increasing concentrations of Vismodegib and subjected to sphere formation assay. Data refers at means ± SEM from two independent experiments. ** $p<0.01$, *** $p<0.001$.

**Supplementary Figure 3**

(A) Frequency and type of MGP mutations in TCGA-HGSOC patients. Data from www.cbioportal.org. (B) Survival analysis of TCGA-HGSOC patients stratified by MGP amplification status (see methods). AMP = amplified MGP, NO AMP = no amplified MGP. Left and right panels show overall survival (OS) and disease-free survival (DFS), respectively. Log-rank p-values are displayed together with hazard ratio (HR, 95% Confidence Interval) and number of patients.

**Supplementary videos**

Time lapse microscopy on the mesothelial clearance assays from MeT5A-RFP (red) co-cultured with OVCAR3 (Suppl. video 1) or COV318 (Suppl. video 2) genetically manipulated for MGP expression. Cultures were monitored for 48 hours with images taken every 2 hours. The spheroids spread, invade, and force the mesothelial cells aside creating a hole in the monolayer. Scale bar = 100 μm.

### Supplementary Figure 1

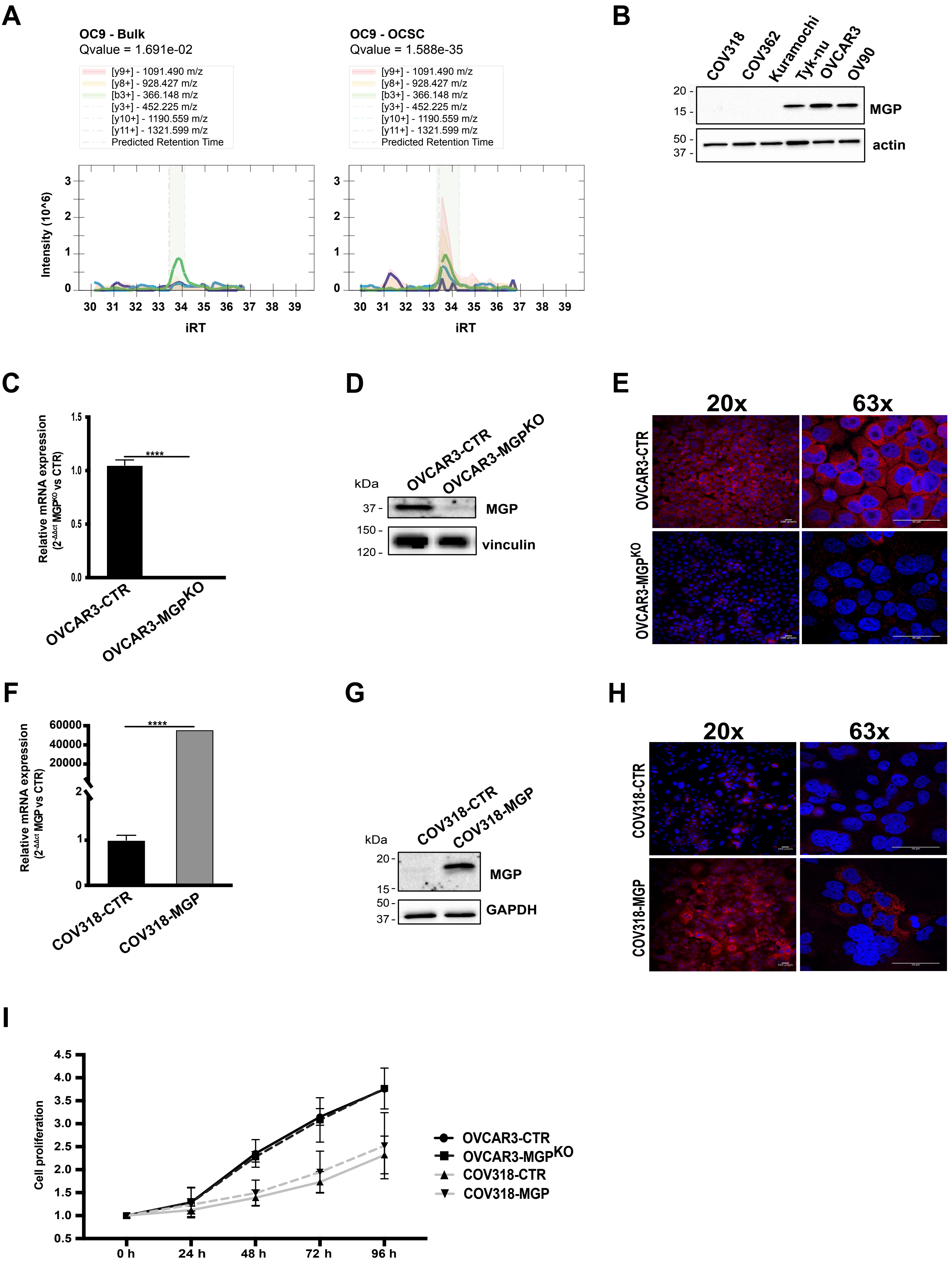

### Supplementary Figure 2

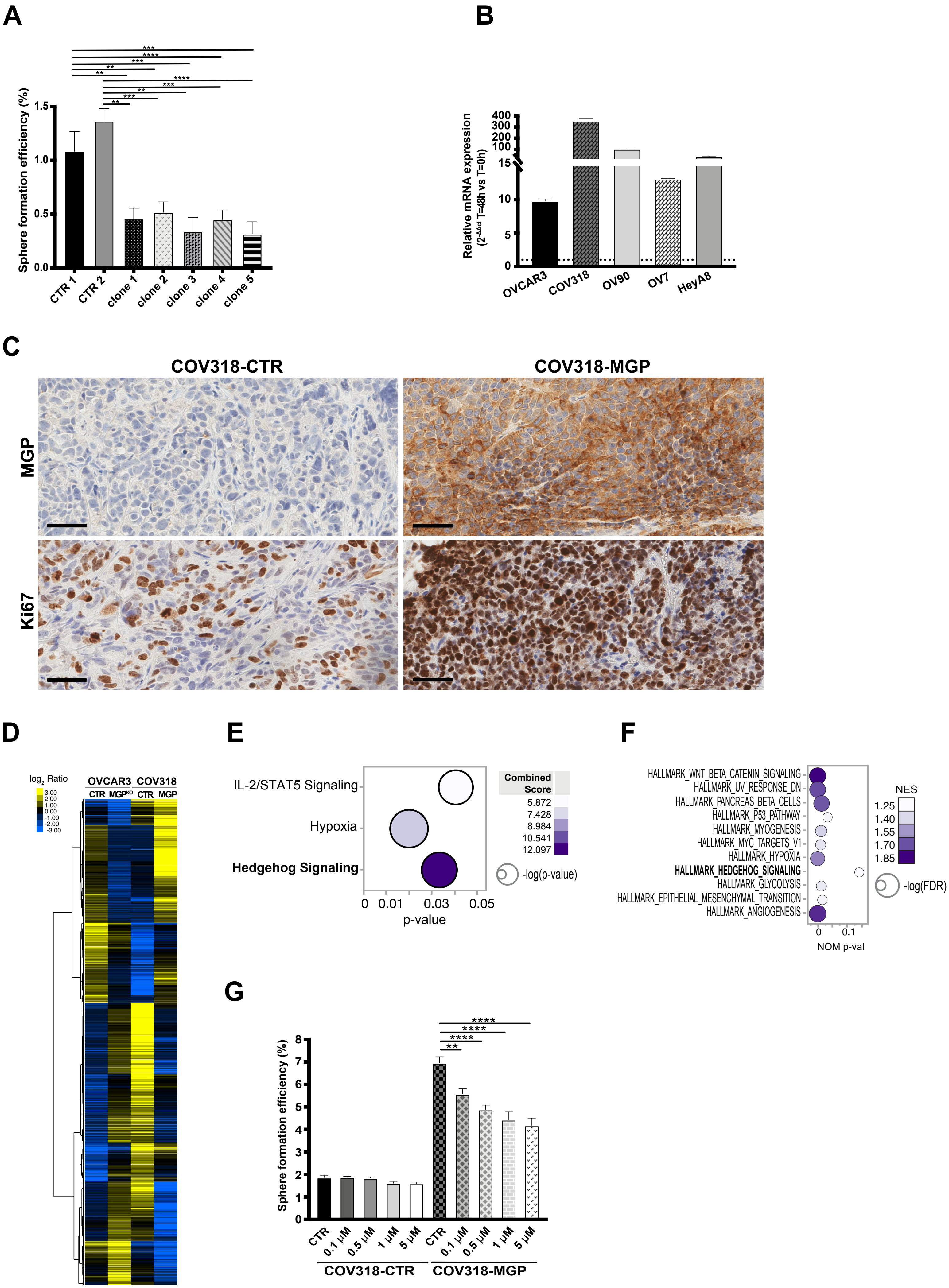

### Supplementary Figure 3

A

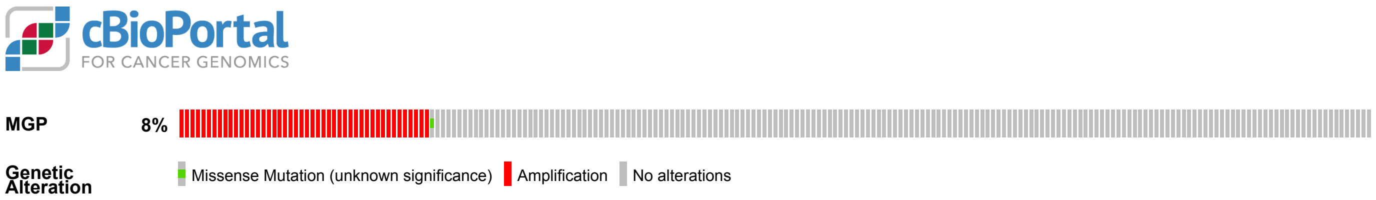


B

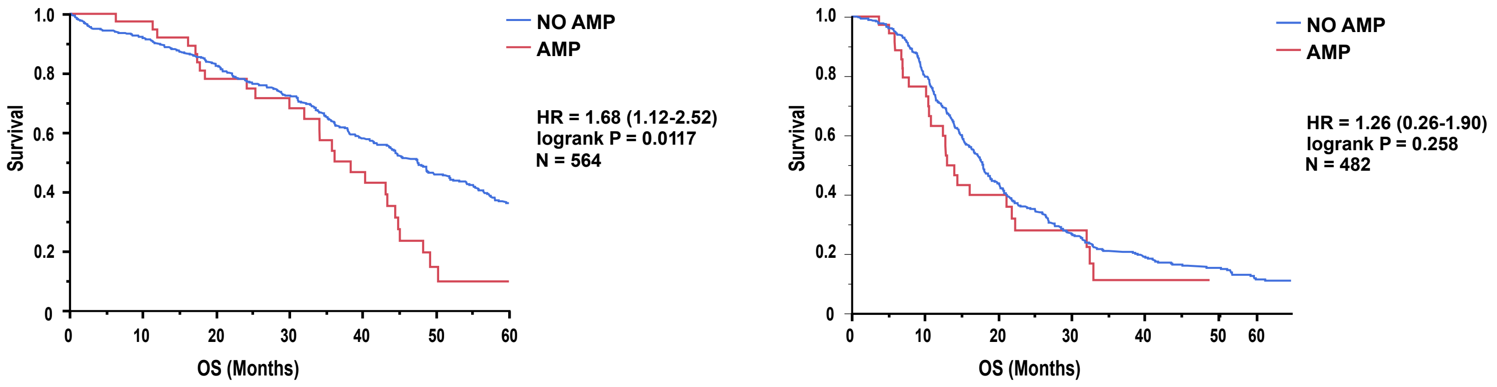
